# Sensitive and Selective Next-Generation FRET-based PKA Biosensors

**DOI:** 10.64898/2026.02.06.702976

**Authors:** Jin-Fan Zhang, Wei Lin, Leyi Huang, Su Hyun Kim, Xinchang He, Michelle S. Frei, Haining Zhong, Sohum Mehta, Jin Zhang

**Author notes:** Corresponding author: Jin Zhang; Sohum Mehta. These authors contributed equally: Jin-Fan Zhang and Wei Lin.

## Abstract

The cyclic AMP (cAMP)/protein kinase A (PKA) signaling pathway regulates diverse cellular processes through precise spatiotemporal control across subcellular compartments. Förster resonance energy transfer (FRET)-based A-kinase activity reporters (AKARs) have enabled live-cell visualization of PKA activity, but their limited dynamic range constrains detection of subtle or compartment-specific signaling events. Here, we present a suite of sixth-generation cyan/yellow FRET-based PKA sensors (the AKAR6 series) with substantially enhanced sensitivity and improved selectivity. Systematic optimization of the FRET donor–acceptor pair and sensor backbone yields superior performance versus previous best-in-class FRET-based AKARs, enabling robust detection of subtle PKA activity changes across diverse experimental modalities, including flow cytometry, fluorescence lifetime-based FRET, and two-photon imaging in brain slices. We further leverage kinome atlas data to engineer a variant with improved selectivity for more accurate visualization of nuclear PKA activity. Using the AKAR6 toolkit, we showed that in contrast to strong GPCR-induced PKA activities across all tested compartments in PC12 cells, growth factors stimulated a significant PKA activity at the trans-Golgi network but no detectable activity at cis-Golgi, signifying highly compartmentalized PKA signaling at the sub-organelle level. Furthermore, NGF and EGF induced sustained and transient PKA activity, respectively, across various intracellular compartments, including the nucleus, suggesting that growth factor-specific temporal controls are maintained across subcellular compartments. Together, the AKAR6 toolkit provides a sensitive, selective, and versatile platform for dissecting compartmentalized PKA signaling across cells and tissues.

## Introduction

The cyclic AMP (cAMP)/protein kinase A (PKA) pathway is one of the most versatile and evolutionarily conserved signaling systems, regulating a broad range of biological processes in different cells and tissues by integrating distinct signals from various extracellular cues such as hormones, neurotransmitters, and growth factors^1-3^. cAMP/PKA signals are transduced across diverse subcellular compartments to achieve precise control of cellular processes. For example, Golgi-localized PKA rapidly translates extracellular signals into structural remodeling of the Golgi and altered protein trafficking.^4^ Cells achieve such exquisite functional specificity through intricate spatiotemporal regulation at the level of PKA via recruitment to A-kinase anchoring proteins (AKAPs)^5^, as well as via both local cAMP degradation by phosphodiesterases (PDEs)^6, 7^ and local cAMP production mediated by intracellular receptor pools, including organelle-specific G protein-coupled receptor (GPCR) signaling from endosomes and the Golgi apparatus.^8, 9^ Deciphering the functional impact of these mechanisms requires advanced tools capable of mapping PKA signaling with high spatial and temporal precision.

Genetically encoded biosensors based on Förster resonance energy transfer (FRET) have transformed our ability to visualize dynamic signaling events in real time within living cells. FRET-based A-kinase activity reporters (AKARs) in particular remain the most widely used tools for monitoring PKA activity. These biosensors consist of a PKA substrate peptide and phosphoamino acid-binding domain (PAABD) sandwiched between two fluorescent proteins (FPs). This modular design has facilitated systematic engineering—such as optimizing the FRET pair, substrate, linker, and PAABD—to steadily improve performance^10, 11^, expand spectral compatibility^12^, and enable alternative readouts such as fluorescence lifetime^13, 14^ or anisotropy imaging^15^, yielding an increasingly versatile AKAR toolkit. Nonetheless, current FRET-based AKARs, even the classic cyan-yellow (C/Y) FRET PKA sensors containing the most widely utilized FRET pair of CFP and YFP, still remain limited by modest dynamic range, limiting their application to detect subtle or compartment-specific signaling events, or challenging conditions such as in vivo imaging in brain tissue. Therefore, there is a continued need to develop enhanced AKAR variants with improved dynamic range and sensitivity.

Here, we report the development of a suite of C/Y FRET-based AKARs with enhanced sensitivity and specificity. These improved AKARs enabled us to robustly detect subtle changes in subcellular PKA activity and specifically uncover the nuclear PKA activity which was obscured by non-specific phosphorylation. Using this new AKAR toolkit, we mapped the subcellular PKA activities in PC12 cells in response to EGF and NGF stimulation, revealing distinct spatial and temporal activation patterns across intracellular compartments.

## Results

### Development and characterization of AKAR6

Like all FRET-based sensors, AKAR reversibly switches between ON and OFF conformational states—controlled by PKA-mediated phosphorylation and phosphatase-mediated dephosphorylation (Figure 1a)—where the change in FRET efficiency between these two states determines the maximum sensor response, or dynamic range. We thus set out to develop a C/Y FRET PKA sensor with increased dynamic range by modifying the current-generation sensor AKAR4 to maximize energy transfer in the high-FRET, ON state and minimize basal energy transfer in the low-FRET, OFF state. To maximize FRET efficiency, we first screened several bright, monomeric CFPs, including mCerulean, mCerulean3, mTurquoise2 and mTurquoise2Δ, which have successfully been used in previous biosensor optimization efforts.^16-19^ Hela cells transiently expressing each construct were stimulated with the adenylyl cyclase activator forskolin (Fsk) and the pan-PDE inhibitor 3-isobutyl-1-methyxanthine (IBMX) to induce maximal PKA activity, followed by washout and treatment with the PKA inhibitor H89 to suppress PKA activity. While none of the CFP variants tested showed a higher yellow-over-cyan (Y/C) emission ratio change compared with the original Cerulean donor FP, they all exhibited faster ON and OFF kinetics than AKAR4 (Cerulean-cpVenus), with mCerulean yielding the fastest activation (T_1/2_ ON = 0.77 ± 0.02 min [mean ± s.e.m.], n = 33 cells; *P =* 6.75×10^-12^ vs. AKAR4) and reversal (T_1/2_ OFF = 0.38 ± 0.02 min, n = 33 cells; *P =* 3.31×10^-22^ vs. AKAR4) (Figure S1). Next, we tested each CFP variant paired with tandem cpVenus (tdcpVenus) (Figure 1a), as tandem acceptors have been reported to increase the effective absorption of the acceptor and ease the dipole alignment between the donor-acceptor pair, thereby improving the dynamic range of FRET sensors.^18, 20-22^ Indeed, each CFP variant, except for mTurquoise2Δ, showed a significantly higher Fsk/IBMX-stimulated Y/C ratio change when paired with tdcpVenus. Notably, the Cerulean-tdcpVenus (ΔR/R_0_ = 79.1% ± 1.2%, n = 34 cells) and mCerulean-tdcpVenus (77.9% ± 1.1%, n = 38 cells) pairs performed the best, showing similarly high dynamic ranges (*P =* 0.483) (Figure S1). Incorporating tdcpVenus only marginally impacted sensor ON kinetics compared with cpVenus (Figure S1). However, tdcpVenus yielded much slower OFF kinetics when paired with either Cerulean (T_1/2_ OFF = 3.16 ± 0.19 min, n = 34 cells, and 0.81 ± 0.05 min, n = 40 cells, for Cerulean-tdcpVenus and Cerulean-cpVenus, respectively; *P* = 1.12×10^-16^) or mCerulean (T_1/2_ OFF = 1.94 ± 0.07 min, n = 38 cells, and 0.38 ± 0.02 min, n = 33 cells, for mCerulean-tdcpVenus and mCerulean-cpVenus, respectively; *P* = 8.99×10^-26^, Figure S1). Taking all of these properties into consideration, we selected the mCerulean-cpVenus and mCerulean-tdcpVenus constructs for additional optimization.

**Figure 1.**
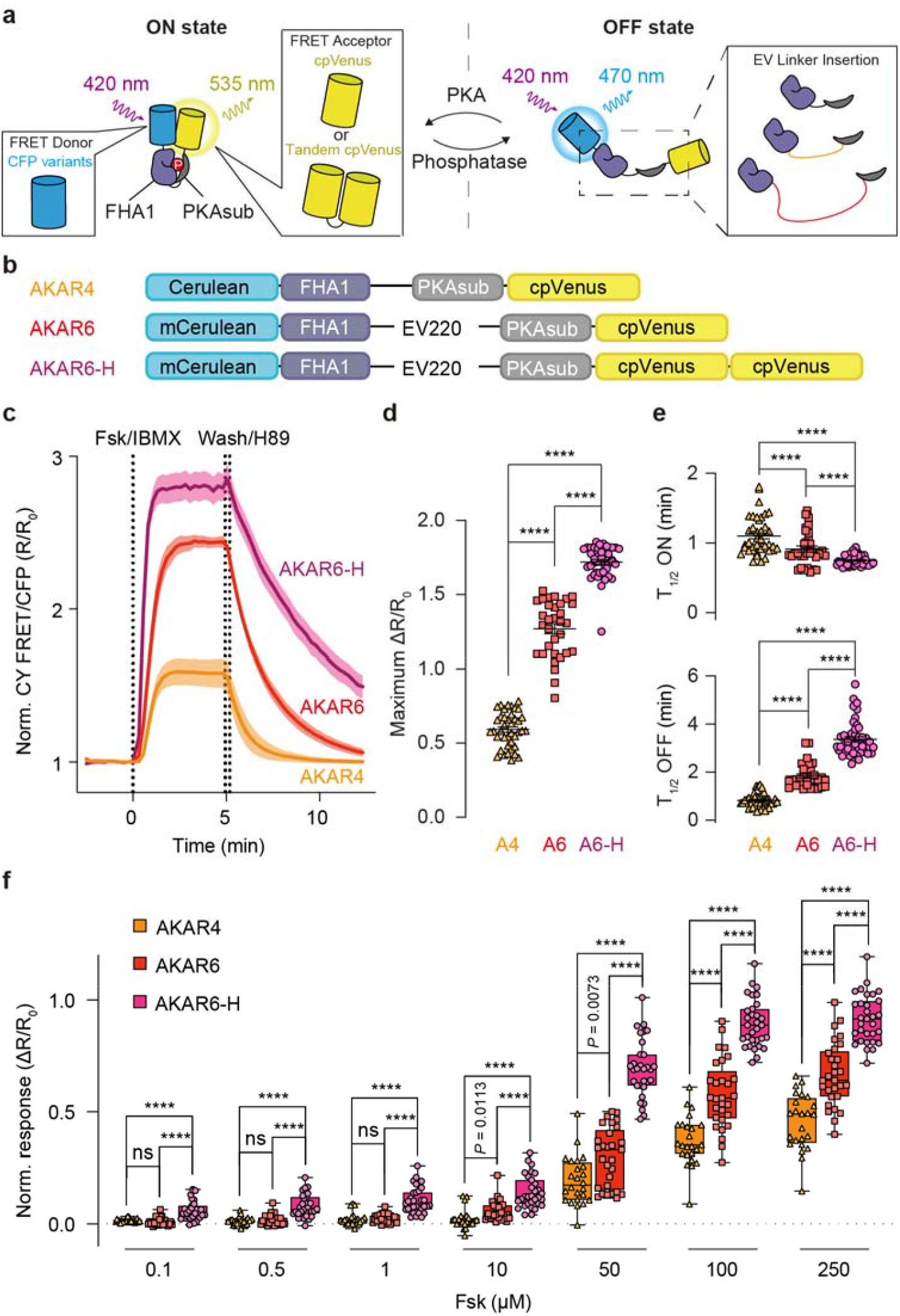
Engineering and characterization of AKAR6 and AKAR6-H. (a) Strategy of optimizing cyan-yellow (CY) FRET-based AKAR. To improve the dynamic range of CY FRET-based AKAR, the FRET efficiency at ON state is enhanced through screening optimal CFP-YFP pairs (left), and FRET efficiency at OFF state is minimized by inserting flexible EV linker (right). (b) Domain structures of AKAR4, AKAR6 and AKAR6-H. (c) Representative average time-course responses in HeLa cells expressing AKAR4 (orange, n = 8 cells), AKAR6 (red, n = 8 cells), AKAR6-H (purple, n = 12 cells) treated with 50 µM Fsk and 100 µM IBMX for 5 min and 20 µM H89 for 10 min. Solid lines indicate mean responses and shaded areas indicate the s.d. (d) Quantification of the maximum Fsk/IBMX-stimulated response of each biosensor in panel c. (e) Quantification of time to half-maximal activation (T_1/2_ ON, upper) and half-maximal inhibition (T_1/2_ OFF, lower) of each biosensor in panel c. AKAR4, n = 40 cells, AKAR6, n = 32 cells, AKAR6-H, n = 44 cells from 3 independent experiments each. Data are mean ± s.e.m. (f) Dose-response (ΔR/R_0_) comparison of AKAR4 (n = 24 cells), AKAR6 (n = 29 cells) and AKAR6-H (n = 31 cells) in HeLa cells. The responses are normalized to Fsk/IBMX treatment. Data are shown as box-and-whisker plots showing the median, interquartile range, min, and max. ns, not significant. \*\*\*\**P* < 0.0001. Data were analyzed using Welch’s ANOVA followed by Dunnett’s test for multiple comparisons.

To further enhance the response amplitude by minimizing OFF-state FRET, we introduced long, flexible Eevee (EV) linkers, consisting of repeating (SAGG)_n_ and (SG)_n_ modules, between the PKA substrate and FHA1 domains (Figure 1a). Previous studies have used EV linkers of different lengths to decrease basal FRET and significantly increase AKAR dynamic range.^11, 23^ Therefore, we systematically screened a panel of EV linkers of increasing length (From 30 to 258 residues) into the mCerulean-cpVenus construct to identify the optimal linker (Figure S2). Increasing EV linker length was inversely correlated with the basal Y/C emission ratio but had no obvious effect on the maximum Fsk/IBMX-induced ratio (Figure S2), consistent with previous reports that longer EV linkers decreased the proximity between PKA substrate and FHA1 domain and increased the proportion of sensors in the OFF state without affecting the ON state.^11^ Calculating the normalized ratio change revealed that the 220-residue EV linker (EV220) yielded the highest overall dynamic range, as well as the fastest ON and OFF kinetics (ΔR/R_0_ = 126.9% ± 3.4%; T_1/2_ ON = 0.91 ± 0.04 min; T_1/2_ OFF = 1.83 ± 0.09 min; n = 32 cells, Figure S3). Further incorporating tdcpVenus into this variant yielded even higher dynamic range (ΔR/R_0_ = 172.0% ± 1.6%, n = 44 cells; *P =* 3.11×10^-19^ vs. cpVenus) and faster ON kinetics (T_1/2_ ON = 0.75 ± 0.01 min, n = 44 cells; *P =* 3.9×10^-^ □ vs. cpVenus) but slower OFF kinetics (T_1/2_ OFF = 3.36 ± 0.11 min, n = 44 cells; *P =* 1.25×10^-16^ vs. cpVenus) (Figure S3). Given the existence of the lifetime-optimized sensor AKAR5^24^, we therefore designated the mCerulean-EV220-cpVenus and mCerulean-EV220-tdcpVenus constructs as AKAR6 and AKAR6-H, respectively (Figure 1b-e). Consistent with their 2.1- and 2.9-fold higher dynamic ranges vs. AKAR4, both AKAR6 and AKAR6-H responded to lower doses of Fsk stimulation than AKAR4 when expressed in HeLa cells (Figure 1f), highlighting their improved sensitivity to detect subtle changes in PKA activity.

### AKAR6 supports multiple application modalities

Given their greatly improved dynamic range and sensitivity, we investigated whether the AKAR6 series is capable of effectively monitoring PKA activity in live cells, regardless of the detection modality, to meet the various research requirements. Flow cytometry is best suited for analyzing single-cell responses in a large population of cells. However, fluorescent biosensors have not been widely used in conjunction with flow cytometry and it is not clear whether temporal dynamics of biosensor responses could be tracked with this modality. Therefore, we sought to monitor AKAR6 emission ratio changes in transiently transfected HEK293T cells via flow cytometry to test the feasibility of this approach. Using time-course flow cytometry to record real-time responses, and applying fluorescence compensation to correct for spectral overlap between CFP, YFP and CY FRET emission, we observed the AKAR6 Y/C emission ratio to increase from 0.26 ± 0.01 to 0.98 ± 0.002 upon stimulation with Fsk/IBMX and then return to baseline upon H89 addition, corresponding to a ∼2.8-fold dynamic range (Figure 2a and S4). These results suggest that AKAR6 can be readily used to analyze single-cell responses with flow cytometry, including via fluorescence activated cell sorting (FACS).

**Figure 2.**
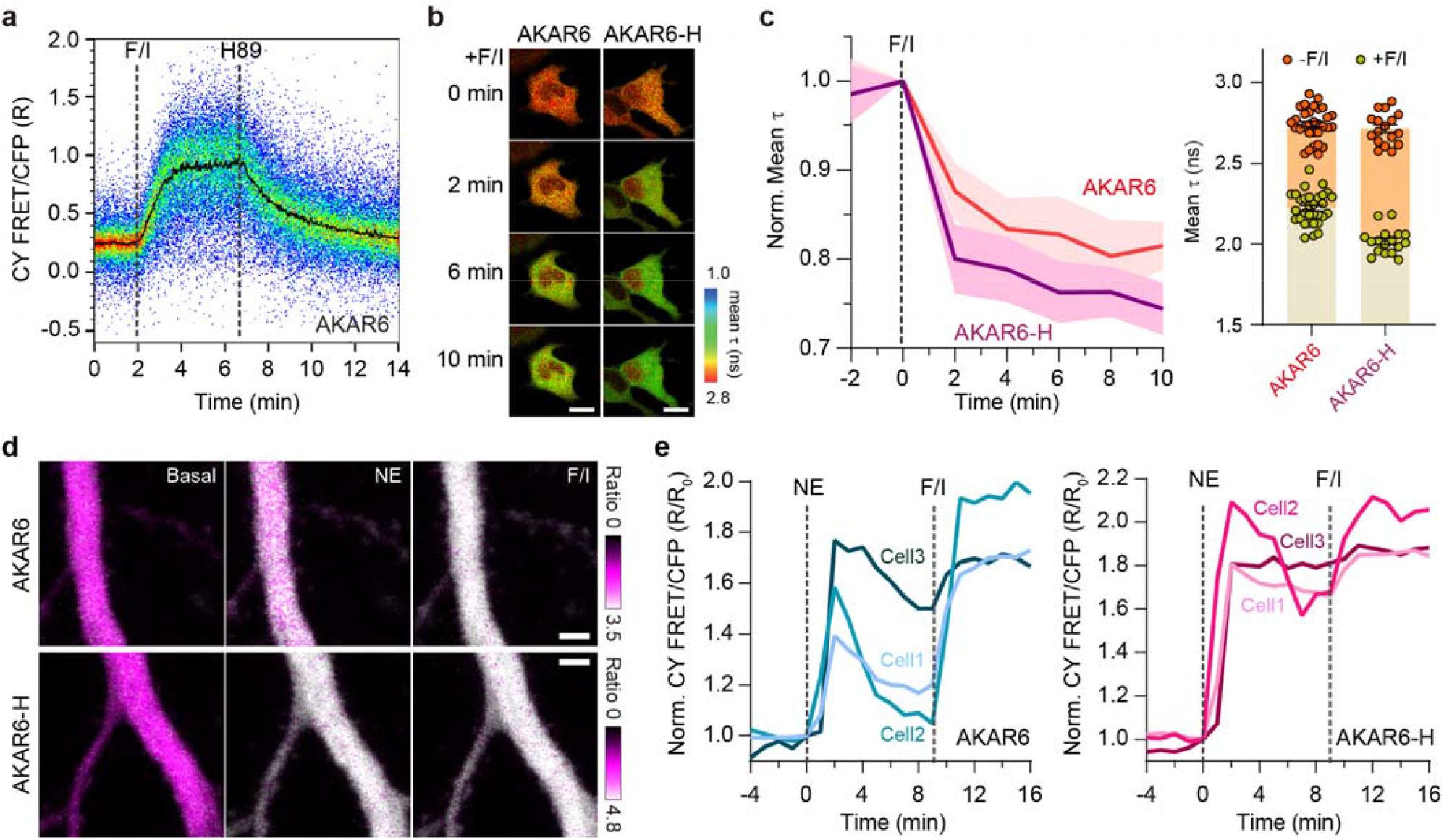
AKAR6 variants enable multi-modal detection of live-cell PKA activity. (a) Flow cytometric measurement of PKA activity in HEK293T cells expressing AKAR6. The scatter plots shows a time-course of the CY FRET/CFP emission ratio in transfected cells, during which Fsk/IBMX and H89 were added at 2 min and 7 min, as indicated. The black curve indicates the average response. Data are representative of three independent experiments (see also Figure S4). (b-c) FLIM-FRET imaging of PKA activity in HeLa cells expressing AKAR6 and AKAR6-H. (b)Representative pseudocolor images illustrate the gradually decreasing donor fluorescence lifetime, corresponding to increasing PKA activity, after Fsk/IBMX stimulation. Scale bar, 10 µm. (c) Left: Representative average time courses showing normalized mean mCerulean lifetime (τ) in individual cells under Fsk/IBMX treatment. AKAR6, n = 12 cells; AKAR6-H, n = 12 cells. Solid lines indicate mean responses and shaded areas indicate s.d. Representative of 3 indepenent experiments. Right: Quantification mean τ before (-) and after (+) Fsk/IBMX treatment. AKAR6, n = 31 cells; AKAR6-H, n = 17 cells; pooled from 3 independent experiments. Data are mean ± s.e.m. (d-e) Representative two-photon ratiometric images (d) and response traces (e) from AKAR6 and AKAR6-H expressed in acute brain slices and treated with the indicated stimuli (n = 3 neurons from 3 brain slices each).

In addition to intensity-based ratiometric imaging, FRET-based biosensors enable fluorescence lifetime imaging microscopy (FLIM), since the fluorescence lifetime of the FRET donor will shorten when intramolecular FRET occurs. Lifetime-based PKA biosensors such as AKAR5^24^ have been developed containing mEGFP and sREACh as the FRET pair. Although AKAR5 exhibits robust fluorescence lifetime changes in response to maximal PKA stimulation, the dynamic range was less than 0.2 ns. We therefore tested whether the enhanced performance of AKAR6 extends to FLIM-FRET imaging by measuring lifetime changes in HEK293T cells expressing AKAR6 sensors. Consistent with our intensity-based imaging results, AKAR6-H showed the largest response, with the fluorescence lifetime (τ) of the mCerulean donor decreasing from 2.71 ± 0.02 ns to 2.01 ± 0.02 ns (Δτ = 0.7 ns, ∼25.7% normalized response, n = 17 cells) in response to PKA stimulation by Fsk/IBMX treatment, whereas AKAR6 showed a more modest lifetime change from 2.74 ± 0.02 ns to 2.22 ± 0.02 ns (Δτ = 0.52 ns, ∼18.9% normalized response, n = 31 cells, Figure 2b,c). Nevertheless, both sensors outperformed the 0.2 ns lifetime change reported for AKAR5, indicating that AKAR6 variants offer enhanced performance for fluorescence lifetime imaging applications.

Lastly, we tested whether AKAR6 sensors are sensitive enough for tissue imaging. We introduced AKAR6 and AKAR6-H into brain slices using the biolistic method and then performed 2-photon imaging to monitor the Y/C emission ratio under 850-nm excitation (Figure S5). Using time-course imaging, we monitored PKA activity changes in the apical dendrites of CA1 neurons in cultured hippocampal slices treated with 1 µM norepinephrine (NE) followed by Fsk/IBMX (Figure 2d). According to the sensors’ response traces obtained from individual neurons, AKAR6-H exhibited strong responses to the physiological ligand NE, which were comparable to those triggered by Fsk/IBMX stimulation (Figure 2e). Although somewhat smaller in amplitude compared with AKAR6-H, the AKAR6 responses more clearly resolved the transient dynamics of NE-stimulated PKA activity (Figure 2e), highlighting distinct advantages of each sensor for deep tissue-imaging applications. Overall, these results suggest that these next-generation AKARs provide high performance and sensitivity to enable versatile application modalities.

### Engineering of AKAR6-S with improved kinase selectivity

Given the enhanced sensitivity of AKAR6, we next set out to investigate the spatial regulation of PKA activity in PC12 cells, which are known to undergo proliferation and neuronal differentiation in response to distinct signaling dynamics induced by epidermal (EGF) or nerve growth factor (NGF), respectively.^25^ Previously, we observed EGF and NGF stimulation to trigger PKA activity with distinct temporal signatures at the plasma membrane in PC12 cells using AKAR4, but were unable to detect cytosolic PKA activation.^26^ In contrast, PC12 cells expressing cytosolic AKAR6 (AKAR6-NES) showed a clear NGF-stimulated response (ΔR/R_0_ = 5.9% ± 0.7%, n = 15 cells) (Figure S6), which was abolished in cells expressing a potent and specific PKA inhibitor peptide (PKI-mCherry-NES) (Figure S6b, c), highlighting the sensitivity of AKAR6. We also observed a strong NGF-induced response from nuclear AKAR6 (AKAR6-NLS), yet co-expressing nuclear PKI (PKI-mCherry-NLS) had no effect despite fully suppressing the Fsk/IBMX-induced AKAR6-NLS response (Figure S6e,f), suggesting the NGF-induced response is PKA independent. Intriguingly, treating AKAR6-NLS and PKI-mCherry-NLS co-expressing PC12 cells with the ATP-competitive inhibitor H89 completely reversed the NGF-induced response (ΔR_inh_/ΔR_max_ = −131.7% ± 7.3%, n = 20 cells, Figure S7a). Despite its widespread use as a PKA inhibitor, H89 is noted to act on numerous other targets in the literature,^27^ and among at least eight other kinases inhibited by H89, we identified mitogen- and stress-activated kinase 1 (MSK1) as the most likely candidate responsible for the NGF-induced AKAR6-NLS response. MSK1 and PKA not only share strikingly similar consensus phosphorylation sequences^28^, but MSK1 is a known nuclear target downstream of the NGF/TrkA-ERK pathway.^29^ Indeed, blocking successive nodes of the TrkA-ERK-MSK1 pathway by treating NGF-stimulated PC12 cells with MEK, ERK and MSK1 inhibitors was able to reverse the NGF-induced AKAR6-NLS response (ΔR_inh_/ΔR_max_ within 10 min, U0126 (to MEK), −58.0% ± 8.3%, n = 14 cells; SCH772984 (to ERK), −52.5% ± 5.1%, n = 25 cells; SB747651A (to MSK1), −103.6% ± 2.5%, n = 19 cells) (Figure S7b-e). Furthermore, purified AKAR6 protein was robustly phosphorylated *in vitro* following incubation with either purified PKA catalytic subunit or a hyperactive variant of human MSK1, dMSK^30^, as indicated by a shift in the emission spectrum (Figure 3a,b).

**Figure 3.**
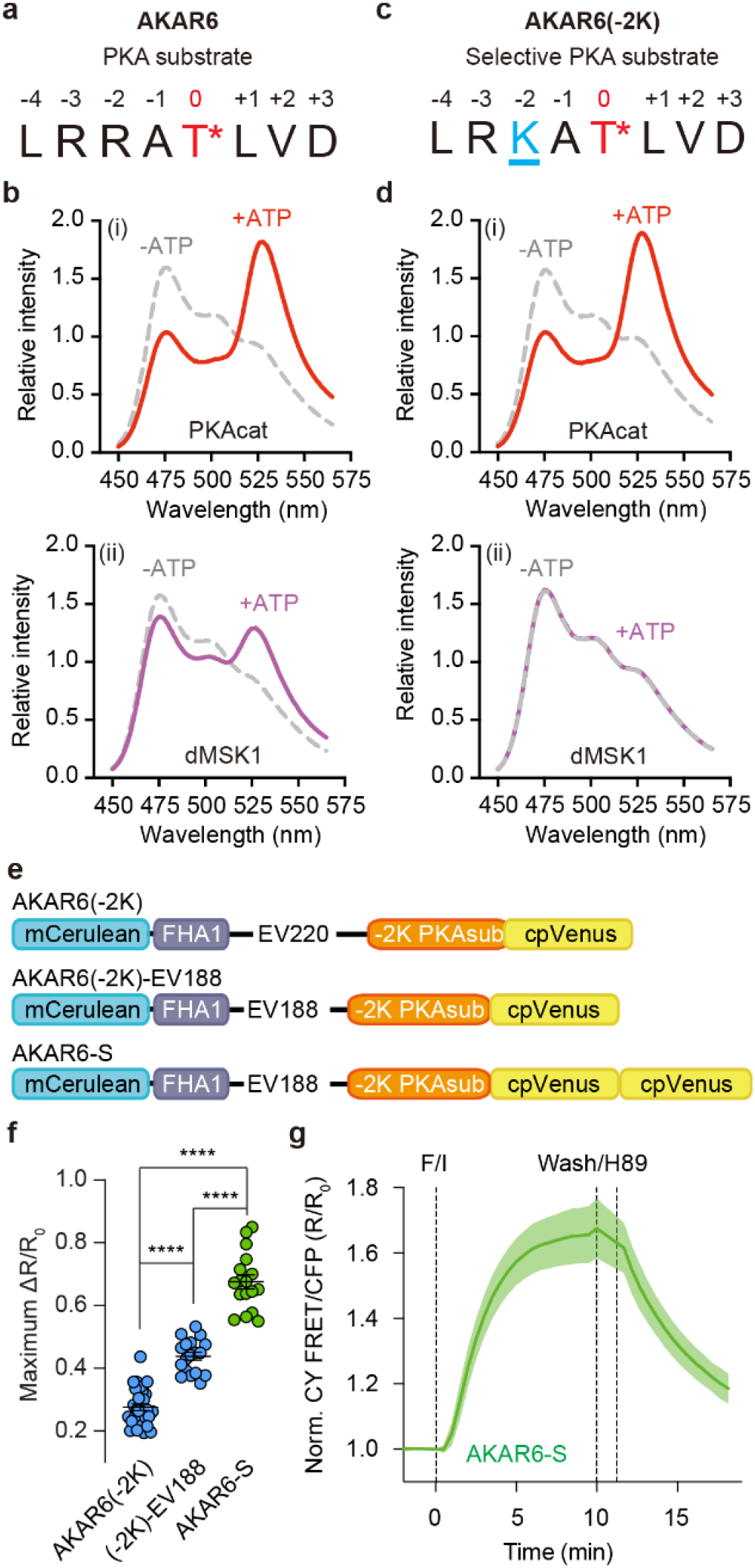
Engineering and characterization of selective AKAR6-S. (a-b) Amino acid sequence of original PKA substrate (a) and AKAR6 emission spectra collected under 420 nm excitation *in vitro* (b) following incubation with PKA catalytic subunit (left) or truncated human MSK1 (right) in the presence (+, solid curves) or absence (-, dashed curves) of ATP. (c-d) Amino acid sequence of re-engineered PKA substrate with −2K mutation (c) and *in vitro* emission spectra of AKAR6(−2K) collected under 420 nm excitation (d) following incubation with PKA catalytic subunit (left) or truncated human MSK1 (right) in the presence (+, solid curves) or absence (-, dashed curves) of ATP. (e) Domain structures of AKAR6(−2K), AKAR6(−2K)-EV188 and AKAR6-S. (f) Quantification of maximum Fsk/IBMX-stimulated responses (from left to right: n = 30, 18, and 17 cells). \*\*\*\**P* < 0.0001. ns, not significant. Data were analyzed using Welch’s ANOVA followed by Dunnett’s test for multiple comparisons. Data are mean ± s.e.m. (g) Average time-course responses in HeLa cells expressing AKAR6-S (n = 17 cells). Solid lines indicate mean responses and shaded areas indicate the s.d. Data from 3 independent experiments.

Our data reveal that the PKA substrate long relied on for AKAR development can also be phosphorylated by nuclear MSK1. We therefore set out to improve the selectivity of AKAR6 by screening a panel of PKA substrate peptides containing amino acid substitutions intended to reduce MSK1 sensitivity while retaining PKA phosphorylation, guided by a recent Ser/Thr-kinome-wide substrate atlas^28^. When tested in both AKAR6 and AKAR6-NLS backgrounds, only the modified substrate carrying an Arg-to-Lys substitution at the −2 position (LR<u>K</u>ATLVD; −2K) yielded sensors that responded to Fsk/IBMX stimulation in HeLa cells (ΔR/R_0_, −2K v.s. T/A 27.6% ± 1.1%, n = 30 cells v.s. 0.5% ± 0.1%, n = 27 cells, *P* = 1.11×10^-20^) (Figure S8a) and showed no response to NGF stimulation in PKI-mCherry-NLS-expressing PC12 cells (ΔR/R_0_, 1.9% ± 0.6%, n = 16 cells v.s. 0.7% ± 0.1%, n = 17 cells, *P =* 0.4492) (Figure S8b). Consistent with these results, purified recombinant AKAR6(−2K) was only phosphorylated by PKA but not MSK1 *in vitro* (Figure 3c,d). A similar effect was observed with ExRai-AKAR2, which uses the same substrate as AKAR6 but has a different fluorescence readout, as purified ExRai-AKAR2(−2K) was selectively phosphorylated by PKA (Figure S9a), and ExRai-AKAR2-2K-NLS showed no response to NGF stimulation in PC12 cells co-expressing PKI-mCherry-NLS (ExRai-AKAR2[−2K]-NLS, ΔR/R_0_ = 5.8% ± 1.8%, n = 12 cells; ExRai-AKAR2-NLS, ΔR/R_0_ = 38.7% ± 6.0%, n = 12 cells; *P =* 6.64×10^-5^) (Figure S9b).

Importantly, AKAR6(−2K) exhibited a weak dynamic range (ΔR/R_0_ = 27.6% ± 1.1%, n = 30 cells) in response to Fsk/IBMX in HeLa cells (Figure S10). Revisiting our original AKAR6 optimization strategies, we found that switching to a slightly shorter EV linker (EV188; ΔR/R_0_ = 43.8% ± 1.3%, n = 18 cells) and replacing cpVenus with tdcpVenus (ΔR/R_0_, = 67.5% ± 2.2%, n = 17 cells) produced the greatest improvements in dynamic range (Figure 3e-g), while also maintaining relatively fast kinetics among all variants (Figure S10). This sensor, designated AKAR6-S, also showed no response to stimulation of other Ser/Thr kinases, including protein kinase C (PKC), calcium–calmodulin-dependent protein kinase II (CaMKII), adenosine monophosphate-activated kinase (AMPK) and Akt (Figure S11). Thus, while AKAR6 remains suitable for most studies of PKA signaling, AKAR6-S represents an important addition for applications where greater selectivity is particularly critical, such as the nucleus.

### Mapping subcellular PKA activity dynamics in PC12 cells

We previously observed PKA activity with distinct temporal signatures at the plasma membrane in response to NGF and EGF signaling in PC12 cells. However, due to the limited sensitivity and selectivity of previous AKARs, whether growth factor-stimulated PKA signaling propagates to other subcellular compartments remains unclear. To address this gap, we generated subcellular targeted AKAR6 or AKAR6-S variants to localize them to plasma membrane, bulk cytosol, trans-Golgi network, cis-Golgi, endoplasmic reticulum (ER), mitochondria and nucleus in PC12 cells (Figure 4a and S12). Each sensor showed good targeting, as confirmed by strong colocalization with the corresponding localization marker, and responded strongly to maximal PKA stimulation with Fsk/IBMX (ΔR/R_0_, PM-AKAR6, 23.3% ± 2.7%, n = 13, AKAR6-NES, 75.2% ± 4.0%, n = 13, Trans-Golgi-AKAR6, 61.6% ± 3.4%, n = 19, Cis-Golgi-AKAR6, 75.3% ± 3.1%, ER-AKAR6, 38.4% ± 2.2%, n = 24, Mito-AKAR6, 35.6% ± 2.0%, n = 17, AKAR6-S-NLS, 10.7% ± 0.6%, n = 14, Figure 4b). We also observed widespread and sustained PKA activity at each subcellular compartment in cells treated with 100 nM pituitary adenylate cyclase activating polypeptide (PACAP) (Figure 4c), an adrenomedullary neurotransmitter that induces PC12 cell neuronal differentiation and neurite outgrowth via both PKA-dependent and independent (e.g., Rap1-ERK) cAMP signaling^31^ through activation of the Gα_s_-linked GPCR PAC1^32^. Normalizing the responses with respect to Fsk/IBMX treatment revealed that PACAP stimulation triggered maximal or near-maximal PKA activity at each subcellular location (ΔR/ΔR_max_: PM-AKAR6, 102.7% ± 1.1%, n = 14; AKAR6-NES, 100.5% ± 1.6%, n = 15; Trans-Golgi-AKAR6, 100.0% ± 0.7%, n = 18; Cis-Golgi-AKAR6, 95.1% ± 1.1%, n = 18; ER-AKAR6, 102.5% ± 1.1%, n = 15; Mito-AKAR6, 99.3% ± 0.9%, n = 10; AKAR6-S-NLS, 78.3% ± 4.2%, n = 8) (Figure 4f), suggesting that PACAP-induced PAC1 activation leads to robust and sustained PKA signaling throughout the PC12 cell interior.

**Figure 4.**
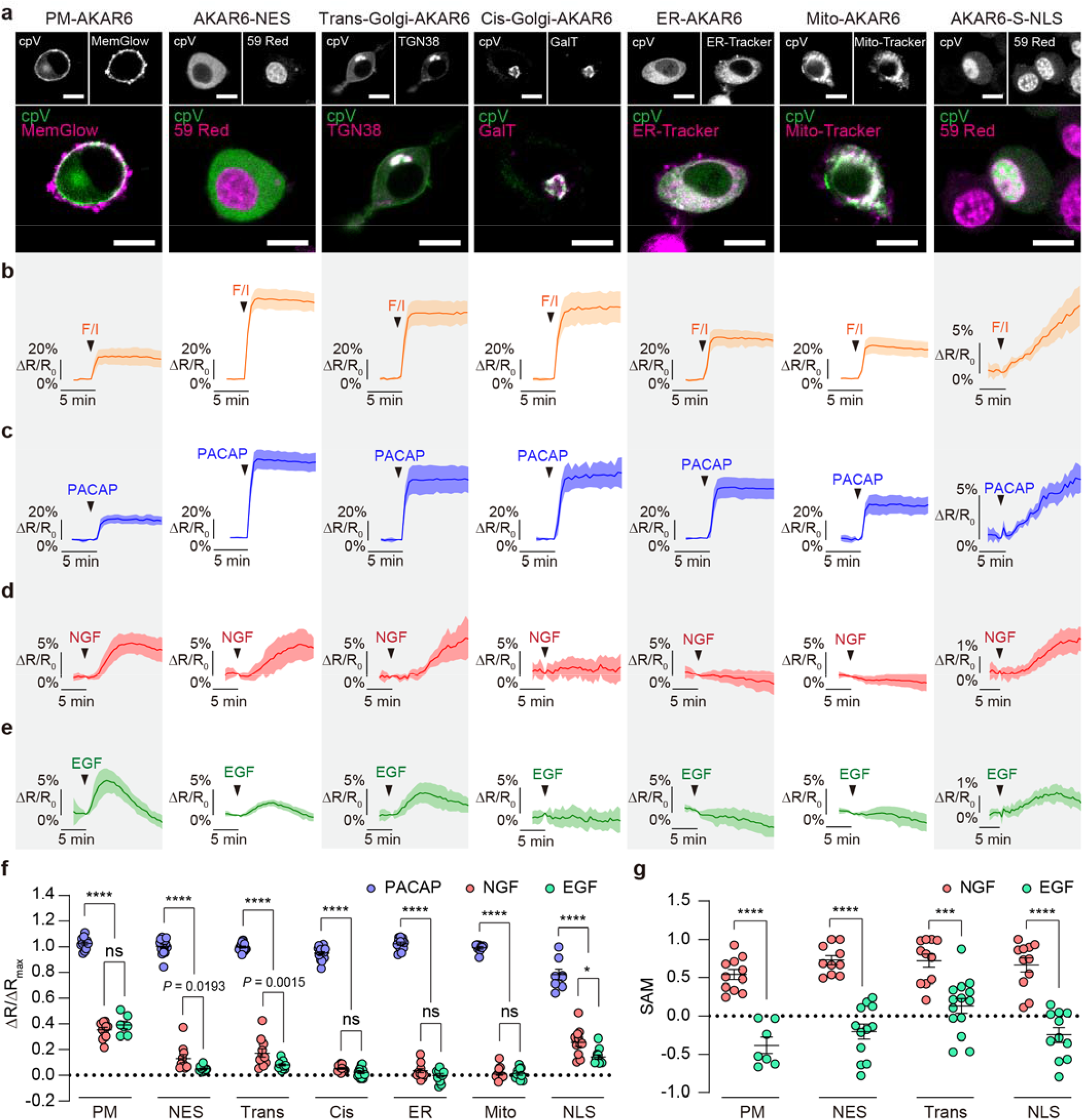
AKAR6 and AKAR6-S allow robust detection of subcellular PKA activities. (a) Representative images of PC12 cells expressing PM-AKAR6, AKAR6-NES, Trans-Golgi-AKAR6, Cis-Golgi-AKAR6, ER-AKAR6, Mito-AKAR6 and AKAR6-S-NLS (YFP channel;green in overlay) along with the indicated localization markers (magenta in overlay): plasma membrane, MemGlow 640; trans-Golgi, TGN38-mCherry; cis-Golgi, GalT-mCherry; ER, ER-Tracker red; mitochondria, Mito-Tracker red; nucleus, SYTO 59 red. Images are representative of 3 independent experiments per condition. Scale bars, 10 □ μm. (b-e) Average time-course responses of subcellularly targeted AKAR6 and AKAR6-S in PC12 cells treated with Fsk/IBMX (b, orange), 100 nM PACAP (c, blue), 200 ng/mL NGF (d, red) or 100 ng/mL EGF (e, green). Solid lines indicate mean responses; shaded areas indicate s.d. (PM-AKAR6, F/I, n = 5 cells, PACAP, n = 5 cells, NGF, n = 11 cells, EGF, n = 7 cells; AKAR6-NES, F/I, n = 5 cells, PACAP, n = 6 cells, NGF, n = 10 cells, EGF, n = 12 cells; Trans-Golgi-AKAR6, F/I, n = 7 cells, PACAP, n = 9 cells, NGF, n = 11 cells, EGF, n = 14 cells; Cis-Golgi-AKAR6, F/I, n = 11 cells, PACAP, n = 6 cells, NGF, n = 13 cells, EGF, n = 17 cells; ER-AKAR6, F/I, n = 5 cells, PACAP, n = 6 cells, NGF, n = 12 cells, EGF, n = 12 cells; Mito-AKAR6, F/I, n = 6 cells, PACAP, n = 7 cells, NGF, n = 17 cells, EGF, n = 12 cells; AKAR6-S-NLS, F/I, n = 6 cells, PACAP, n = 4 cells, NGF, n = 11 cells, EGF, n = 12 cells). (f) Maximum responses of subcellularly targeted AKAR6 and AKAR6-S in PC12 cells treated with PACAP, NGF or EGF normalized to Fsk/IBMX (ΔR/ΔR_max_). (PM-AKAR6, n = 14, 11 and 7 cells; AKAR6-NES, n = 15, 10 and 12 cells; Trans-Golgi-AKAR6, n = 18, 11 and 14 cells; Cis-Golgi-AKAR6, n = 18, 13 and 17 cells; ER-AKAR6, n = 12, 13 and 12 cells; Mito-AKAR6, n = 10, 17 and 12 cells; AKAR6-S-NLS, n = 8, 11 and 12 cells). \*\*\*\**P* < 0.0001. ns, not significant. Data were analyzed using ordinary one-way ANOVA followed by Dunnett’s multiple comparison test. (g) The comparison of NGF- and EGF-induced response dynamics of subcellularly targeted AKAR6 and AKAR6-S, quantified as the sustained activity metric at 20 minutes (SAM20) for PM-AKAR6 (n = 11 and 7 cells), AKAR6-NES (n = 10 and 12 cells), Trans-Golgi-AKAR6 (n = 11 and 14 cells) and SAM30 for AKAR6-S-NLS (n = 11 and 12 cells). \*\*\*\**P* < 0.0001. Data were analyzed using unpaired two-tailed Student’s *t*-test. Data are mean ± s.e.m.

In contrast to GPCR signaling, we observed distinct compartmentation of PKA responses at different subcellular locations following NGF and EGF stimulation, (Figure 4d-f). Both NGF and EGF induced robust responses at the plasma membrane in PM-AKAR6-expressing PC12 cells (ΔR/ΔR_max_, NGF 35.2% ± 1.9%, n = 11 v.s. EGF 39.0% ± 2.9%, n = 7, *P =* 0.3736). Consistent with our previous observations using AKAR4,^26^ NGF-induced PKA activity was sustained, whereas EGF led to a transient response, as quantified using a sustained activity metric (SAM; see Methods) (SAM[NGF] = 0.54 ± 0.07, n = 11 cells; SAM[EGF] = −0.38 ± 0.10, n = 7 cells; p < 0.0001, Figure 4g and S13). AKAR6 also robustly reported both NGF- and EGF-induced PKA responses when targeted to the cytosol, trans-Golgi network and nucleus (Figure 4d-f, S13), but detected little activity when targeted to the cis-Golgi, ER or mitochondria (Figure. 4d-f, S13), revealing that growth factor-induced PKA activity is highly spatially regulated. The contrast between trans and cis-Golgi suggests highly compartmentalized PKA signaling at the sub-organelle level. The distinct growth factor-induced response dynamics were also preserved intracellularly, with NGF triggering sustained PKA activity and EGF stimulation yielding transient PKA activity at all responding intracellular compartments (SAM: AKAR6-NES, NGF 0.73 ± 0.06, n = 10 cells v.s. EGF −0.20 ± 0.10, n = 12 cells, P < 0.0001; Trans-Golgi-AKAR6, NGF 0.72 ± 0.09, n = 11 cells v.s. EGF 0.13 ± 0.10, n = 14 cells, *P =* 0.0002; AKAR6-S-NLS, NGF 0.67 ± 0.09, n = 11 cells v.s. EGF −0.25 ± 0.09, n = 12 cells, P < 0.0001, Figure 4g). Notably, whereas NGF and EGF induced similar normalized PKA activity levels at the plasma membrane (ΔR/R_max_), the normalized responses from intracellular compartments were lower for EGF stimulation compared with NGF (ΔR/ΔR_max_: AKAR6-NES, NGF 13.0% ± 3.1%, n = 10 v.s. EGF 8.1% ± 0.8%, n = 12, *P =* 0.0193; Trans-Golgi-AKAR6, NGF 17.3% ± 3.3%, n = 11 v.s. EGF 4.8%±0.2%, n = 14, *P =* 0.0015; AKAR6-S-NLS, NGF 25.8% ± 3.2%, n = 11 v.s. EGF 14.2% ± 1.8%, n = 11, *P =* 0.0237) (Figure 4d-f), suggesting that EGF-induced PKA signaling is consistently weaker and more transient than NGF-signaling across different subcellular regions. Altogether, the AKAR6 toolkits depicted a clearer picture of subcellular PKA signaling atlas downstream of GPCR and growth factors with spatiotemporal divergence.

## Discussion

In this study, we have developed the AKAR6 toolkit, a suite of next-generation PKA biosensors with significantly improved performance in terms of both sensitivity and selectivity. Greater sensitivity over the previous best-in-class C/Y FRET PKA sensor, AKAR4, was achieved by optimizing both the FRET donor-acceptor pair and the sensor backbone. The high sensitivity of AKAR6 sensors enabled robust monitoring of PKA activity across multiple application modalities, including flow cytometry and FLIM-FRET imaging, achieving comparable performance in tissue slice imaging to that in cultured cells. With the help of positional scanning peptide array data^28^, we also successfully engineered a more selective variant, AKAR6-S, following the unexpected discovery of off-target AKAR6 phosphorylation in PC12 cells by the growth factor-stimulated nuclear kinase MSK1.

Using this enhanced AKAR toolkit, including the more selective AKAR6-S, allowed us to investigate the spatiotemporal regulation of PKA signaling in response to growth factor stimulation in PC12 cells. For both EGF and NGF, PKA activity was primarily confined to the plasma membrane, cytosol, trans-Golgi network, and nucleus. Notably, the cis- and trans-Golgi networks exhibited drastically different PKA signaling, with robust activity observed at the trans-Golgi versus almost no detectable activity at the cis-Golgi, providing a striking example of sub-organelle compartmentation of PKA signaling. Considering the complex and dynamic relationship between the plasma membrane and trans-Golgi network, including frequent vesicle-mediated exchanges, growth factor-stimulated PKA signaling may be specifically directed to the trans face of the Golgi apparatus in a vesicle trafficking-dependent manner. Our results also showed that the characteristic sustained versus transient PKA activity dynamics induced by NGF and EGF were preserved intracellularly, suggesting these growth factor specific temporal controls are achieved via general circuit wiring^33^ instead of compartment-specific feedback mechanisms.^34^ NGF stimulates sustained PKA activity in the nucleus, suggesting that NGF may exert long-term effects on gene regulation. Furthermore, considering that both PACAP and NGF stimulate neurite extension and neuronal differentiation^26, 35^, and both produce sustained PKA activity but with different amplitude, the duration of PKA signaling may be a critical factor guiding cell fate decisions. However, given the similar phosphorylation motifs and substrates of MSK1 and PKA, including CREB^36^, it will be important to precisely tease apart how each signaling axis (RTK– ERK–MSK1 and RTK–cAMP–PKA) contributes to cellular differentiation and proliferation.^26, 37^

In summary, we have presented a sensitive and specific toolkit of next-generation FRET-based PKA sensors. The demonstrated utility of the AKAR6 suite for monitoring subtle PKA activity changes at the subcellular, cellular, and tissue levels, as well as compatibility with various monitoring technologies, highlights their versatility for illuminating kinase signaling across a multitude of applications.

## Data availability

All data supporting the findings of this study are available upon reasonable request. Source data are provided with this paper.

## Acknowledgments

We thank Jared Lee Johnson and Lewis Cantley for their suggestions on kinase motif characterization. This work is supported by the National Institutes of Health (NIH) grants R35 CA197622, R01 DK073368 and R01 CA262815 (to J.Z.) and RF1 MH130784, RF1 NS133599, and R01 NS127013 (to H.Z.); the American Heart Association Predoctoral Fellowship, Award #834472 (to J.F.Z.); the Swiss National Science Foundation (SNSF) grant P2ELP3_199834 (to M.S.F.). The UCSD Microscopy Core is supported by NIH grants P30 NS047101, S10 OD030505.

## Author contributions

J.F.Z., S.M. and J.Z. conceived the project. J.F.Z. and S.M. constructed AKAR variants. X.H. and M.S.F. generated the EV linker library. J.F.Z. and L.H. generated AKAR substrate variants. J.F.Z. performed *in vitro* phosphorylation experiments. J.F.Z., W.L., S.M. and L.H. performed live-cell imaging in HeLa and PC12 cells. S.H.K. performed flow cytometry. W.L. performed FLIM imaging. H.Z. performed 2-photon imaging in organotypic hippocampal slices. J.F.Z., W.L., S.H.K., H.Z. and S.M. analyzed the data. W.L., S.M. and J.Z. supervised the project and coordinated experiments. J.F.Z., W.L. S.M. and J.Z. wrote the manuscript with input from all authors.

## Method

### Plasmids

All primers used for molecular cloning are listed in Table S1. AKAR donor/acceptor variants were generated as follows. AKAR containing a truncated mTurquoise2 (mTurquoise2Δ) FRET donor was constructed by ligating a *Bam*HI/*Sph*I-digested PCR fragment (primers 1 and 2) encoding mTurquoise2Δ PCR-amplified from EpacS^H18818^ (gift of Kees Jalink, Addgene plasmid # 170349) into *Bam*HI/*Sph*I-digested AKAR4. To incorporate tandem-dimer cpVenusE172 (td-pVenus), we first PCR-amplified one copy of cpVenusE172 from EpacS^H188^ using primers 3 and 4, encoding a 5’ *Sac*I site and 3’ linker sequence. This fragment was used to amplify a second copy of cpVenus using primers 5 and 6, encoding an overlapping linker sequence at the 5’ end and a 3’ *Eco*RI site. Both fragments were then inserted into *Sac*I/*Eco*RI-digested AKAR4 and AKAR-mTurquoise2Δ/cpVenus backbones using the NEBuilder HiFi DNA Assembly Kit (New England Biolabs E2621). For mCerulean3 and full-length mTurquoise2 donors, a *Bam*HI/*Sph*I-digested fragment encoding mCerulean3 from CaNAR2^38^ was ligated into *Bam*HI/*Sph*I-digested AKAR4 and AKAR-Cerulean/tdcpVenus backbones, and full-length mTurquoise2 PCR-amplified (primers 7 and 8) from FRESCA2^39^ (kind gift of Margaret Stratton, University of Massacheusetts) was inserted into the same *Bam*HI/*Sph*I-digested backbones using the NEBuilder HiFi DNA Assembly Kit (New England Biolabs E2621). mCerulean was obtained via site-directed mutagenesis using overlapping primers 9 and 10 to introduce an A206K mutation into Cerulean-FHA1-NES^40^, followed by ligation of *Bam*HI/*Sph*I-digested mCerulean into *Bam*HI/*Sph*I-digested AKAR4 and AKAR-Cerulean/tdcpVenus backbones. All constructs were generated in pRSETb and subcloned into pcDNA3 for mammalian expression.

AKAR-mCerulean/cpVenus incorporating a 116-residue EV linker was generated as follows. The plasmid backbone was PCR-amplified using overlapping primers 11 and 12, complementary to the SV40 polyA signaling within the pcDNA3 backbone, paired with primers 13 and 14, designed to insert a *Kpn*I site after the FHA1 domain and *Bsp*EI site before the PKA substrate, respectively, to yield two backbone fragments. Both PCR fragments were then combined with *Kpn*I/*Bsp*EI-digested EV linker fragment from EKAR4^34^ using the NEBuilder HiFi DNA Assembly Kit (New England Biolabs E2621). The EV linker library was generated by PCR-amplifying short EV linker truncations using primers 15 and 16. The resulting truncations were assembled by random Gibson assembly into *Kpn*I/*Bsp*EI-digested AKAR-mCerulean/EV116/cpVenus. Multiple colonies were tested to obtain constructs spanning different EV linker lengths. Finally, AKAR-mCerulean/EV220/tdcpVenus (AKAR6-H) was constructed by ligating a *Bam*HI/*Sac*I-digested fragment of AKAR6 (AKAR-mCerulean/EV220/cpVenus) and *Sac*I/*Eco*RI-digested fragment encoding tdcpVenus into a *Bam*HI/*Eco*RI-digested pcDNA3 backbone. All plasmids were verified by Sanger sequencing (Azenta).

AKAR6 substrate mutants were generated first via Gibson Assembly using NEBuilder HiFi DNA Assembly Kit (New England Biolabs E2621) using primers 17-27 to introduce mutations in PKA substrate of AKAR6. The yielding constructs with random length of EV Linkers were then digested by *Kpn*I/*Bsp*EI and ligated with *Kpn*I/*Bsp*EI digested EV220 Linker to generate AKAR6 substrate mutant with EV220 linker. AKAR6 −2K with different length of EV Linkers were generated by inserting *Kpn*I/*Bsp*EI digested EV Linker with variable length into the *Kpn*I/*Bsp*EI digested AKAR6-2K construct. AKAR6-S was generated first via Gibson Assembly using NEBuilder HiFi DNA Assembly Kit (New England Biolabs E2621) using primers 17 and 19 to introduce the −2K mutation in PKA substrate of AKAR6-H. The yielding constructs with random length of EV Linker were then digested by *Kpn*I/*Bsp*EI and ligated with *Kpn*I/*Bsp*EI digested EV188 Linker to generate AKAR6-S.

For subcellular targeted variants, AKAR6-Kras and AKAR6-NES were generated by subcloning a *Bam*HI/*Age*I-digested fragment containing the majority of AKAR6 into *Bam*HI/*Age*I-digested backbones of pcDNA3-AKAR4-Kras (addgene #61621) and pcDNA3-AKAR4-NES (addgene #64727), respectively. Trans-Golgi-AKAR6, ER-AKAR6 and Mito-AKAR6 variants were generated by subcloning a *Bam*HI/*Eco*RI-digested fragment containing full-length AKAR6 into *Bam*HI/*Eco*RI-digested backbones containing the N-terminal 35 amino acids from Endothelial nitric oxide synthase (eNOS) (MGNLKSVAQEPGPPCGLGLGLGLGLCGKQGPATPA), the N-terminal 27 amino acids from CYP450 (MDPVVVLGLCLSCLLLLSLWKQSYGGG), and N-terminal 30 amino acid leader sequence from DAKAP1 (MAIQLRSLFPLALPGMALLGWWWFFSRKK), respectively. Cis-Golgi-AKAR6 was generated by subcloning a *Bam*HI/*Eco*RI-digested fragment containing full-length AKAR6 without stop-codon into *Bam*HI/*Eco*RI-digested backbones containing the Giantin fragment (3131-3259). AKAR6-S-NLS was generated first via PCR amplifying the fragment from AKAR6-2K using primers 28 and 29 and AKAR6-NLS using primers 30 and 31. The two fragments were then ligated with *Bsp*EI/*Xba*I digested backbones of AKAR6 to generate mCerulean-EV Linker (random length)-(−2K substrate)-tdcpVenus-NLS. The random length EV Linker was then digested by *Kpn*I/*Bsp*EI and inserted with *Kpn*I/*Bsp*EI digested EV188 Linker to generate AKAR6-S-NLS.

### Biosensor purification and *in vitro* characterization

Polyhistidine-tagged AKAR6, AKAR6-2K, ExRai-AKAR2, ExRai-AKAR2-2K and human dMSK1 in pQTEV were transformed into E. coli BL21 cells and purified by nickel affinity chromatography. Briefly, cells were grown at 37 □ °C to an optical density (OD600 nm) of 0.3– 0.4 and then induced overnight at 21 □ °C with 0.4 □ mM IPTG. The cells were pelleted, resuspended in lysis buffer (50 □ mM Tris, pH □ 7.4, 300 □ mM NaCl) containing 1 mM PMSF and Complete EDTA-free Protease Inhibitor Cocktail (Roche), and lysed by sonication. Following centrifugation at 25,000 g for 30 □ min at 4 □ °C, the clarified lysate was loaded onto an Ni-NTA column, and bound protein was subsequently eluted using an imidazole gradient (10– 200 mM). Eluted fractions were analyzed via SDS–PAGE, pooled and concentrated using Amicon Ultra-15 centrifugal columns (30-kD cut-off, Millipore). Protein concentrations were determined using the Pierce BCA Protein Assay Kit (Thermo Fisher Scientific) on a Spark 20M microplate reader (Tecan).

Excitation and emission spectra were obtained on a PTI QM-400 fluorometer using FelixGX v.4.1.2 software (Horiba). Cyan-yellow FRET emission scans were collected under 420-nm excitation, dual-GFP excitation scans were collected at 530-nm emission. Purified AKAR6, AKAR6-2K, ExRai-AKAR2 or ExRai-AKAR2-2K (1 □ μM) were incubated with 5□μg of PKAcat or human dMSK1 respectively for 30 □ min at 30 □ °C in kinase assay buffer (50 □ mM Tris-HCl, pH □ 7.5, 10 □ mM MgCl2, 0.1 □ mM EDTA, 2 □ mM DTT, 0.01% Brij 35) without ATP for unphosphorylated spectra or with 200□μM ATP for phosphorylated spectra.

### Cell culture and transfection

HeLa and HEK293T cells were cultured in Dulbecco’s modified Eagle medium (DMEM, Gibco) supplemented with 10% (v/v) fetal bovine serum (FBS, Sigma) and 1% (v/v) penicillin-streptomycin (Pen-Strep, Sigma-Aldrich). PC12 cells were cultured in DMEM containing 1 □ g □ l^−1^ of glucose, 10% (v/v) FBS, 5% donor horse serum (DHS, Gibco) and 1% (v/v) Pen-Strep. NIH3T3 cells were cultured in DMEM containing 1 □ g □ l^−1^ glucose and supplemented with 10% (v/v) fetal calf serum (ATCC) and 1% (v/v) Pen-Strep. All cells were maintained at 37 □ °C in a humidified atmosphere of 5% □ CO_2_. For imaging experiments, cells were plated onto sterile 35-mm glass-bottomed dishes and grown to 50–70% confluence. Transient transfection was performed using Lipofectamine 2000 (Invitrogen), after which cells were cultured for an additional 24 □ h (HeLa, HEK293T, NIH3T3) or 48 □ h (PC12). PC12 cells were changed to reduced serum medium (5% FBS, 1% DHS) 24 □ h before imaging. NIH3T3 cells were changed to serum-free DMEM immediately before transfection and serum-starved for 24 □ h before imaging.

### Biosensor localization assay

PC12 cells expressing AKAR6-Kras were stained with MemGlow^TM^ 640 Fluorogenic Membrane Probe (MG04, Cytoskeleton, Inc.) at 100 nM in Hank’s balanced salt solution (HBSS) for 10 min. PC12 cells expressing AKAR6-NES and AKAR6-S-NLS were stained with SYTO™ 59 Red Fluorescent Nucleic Acid Stain (Invitrogen) at 5 µM in HBSS for 30 min. PC12 cells expressing Mito-AKAR6 and ER-AKAR6 were stained with MitoTracker RED (Invitrogen) and ERTracker RED (Invitrogen), respectively, at 1 □ mM in HBSS for 30 min. PC12 cells expressing Trans-Golgi-AKAR6 were co-transfected with TGN38-mCherry. PC12 cells expressing Cis-Golgi-AKAR6 were co-transfected with mCherry-GalT (Addgene # 87327). Co-imaging experiments were performed on Leica SP8 confocal microscope. cpVenus in each AKAR6 variant was exited at 488 nm and collected with a 505–550 nm band-pass filter. MemGlow^TM^ 640 was exited at 633 nm and collected with a 660–700 nm band-pass filter. SYTO™ 59 was exited at 610 nm and collected with a 655–690 nm band-pass filter. ERTracker RED, MitoTracker RED and mCherry were exited at 561 nm and collected with a 590–640 nm band-pass filter. The line profile for AKAR6-Kras, AKAR6-NES and AKAR6-S-NLS were measured using ImageJ, The Pearson’s coefficients for Trans-Golgi-AKAR6, Cis-Golgi-AKAR6, ER-AKAR6 and Mito-AKAR6 were calculated using Coloc 2 mode in ImageJ.^41^

### Time-lapse fluorescence imaging

Cells were washed twice with HBSS and subsequently imaged in HBSS in the dark at room temperature. Forskolin (Fsk) (Calbiochem), IBMX (Sigma), H89 (Sigma), NGF (Harlan Laboratories), EGF (Sigma), PMA (LC Laboratories), Thapsigargin (Thermo), 2-deoxyglucose (Sigma), PDGF (Sigma), SCH772984 (selleckchem), U0126 (Sigma) and SB-747651A (MedChemExpress) were added as indicated.

HeLa, HEK293T, PC12 and NIH3T3 cells were imaged on a Zeiss AxioObserver Z1 microscope (Carl Zeiss) equipped with a Definite Focus system (Carl Zeiss), a 40×/1.3 □ NA oil objective and a Photometrics Evolve 512 EMCCD (Photometrics) and controlled by METAFLUOR 7.7 software (Molecular Devices). CY FRET/CFP emission ratio imaging was performed using a 420DF20 excitation filter, a 455DRLP dichroic mirror and two emission filters (473DF24 for cyan fluorescent protein and 535DF25 for yellow fluorescent protein). Dual GFP excitation-ratio imaging was performed using 480DF30 and 405DF40 excitation filters, a 505DRLP dichroic mirror and a 535DF45 emission filter. RFP intensity was imaged using a 555DF25 excitation filter, a 568DRLP dichroic mirror and a 650DF100 emission filter. All filter sets were alternated using a Lambda 10–2 filter changer (Sutter Instruments). Exposure times for each channel were ranged from 200 to 500 □ ms, and images were acquired every 30 □ s.

Raw fluorescence images were corrected by subtracting the background fluorescence intensity of a cell-free region from the emission intensities of biosensor-expressing cells at each time point. CY FRET/CFP emission ratios (Em535/Em473), GFP excitation ratios (Ex480/Ex405) and RFP, fluorescence intensities were then calculated at each time point. All biosensor response timecourses were subsequently plotted as the normalized fluorescence intensity or ratio change with respect to time zero (ΔF/F_0_ or ΔR/R_0_), calculated as (F □ − □ F_0_)/F_0_ or (R □ − □ R_0_)/R_0_, where F and R are the fluorescence intensity and ratio value at a given time point, and F_0_ and R_0_ are the initial fluorescence intensity or ratio value at time zero, which was defined as the time point immediately preceding drug addition. Maximum intensity (ΔF/F_0_) or ratio (ΔR/R_0_) changes were calculated as (F_max_ □ − □ F_0_)/F_0_ or (R_max_ □ − □ R_0_)/R_0_, where F_max_ and F_0_ or R_max_ and R_0_ are the maximum and starting intensity or ratio value recorded after stimulation, respectively. Graphs were plotted using GraphPad Prism 8 (GraphPad Software).

### Flow cytometry

HEK293T cells (2 × 10 □ cells per well) were seeded into 6-well plates containing 2 mL of complete culture medium and maintained at 37°C in a humidified incubator with 5% COL. After a 24-hour incubation, cells were transfected with 500 ng of plasmid DNA using 3 µL of PolyJet reagent according to the manufacturer’s instructions. Following an additional 24-hour incubation, cells were washed with DPBS and detached using trypsin. Approximately 1 × 10 □ cells were collected by centrifugation, washed again with DPBS, and resuspended in 1 mL of DPBS supplemented with 1% FBS and 1% antibiotic-antimycotic solution (Gibco). Cells were filtered through a cell strainer and maintained on ice until analysis.

Fluorescent signals were measured using LSRFortessa X-20 cell analyzer (BD Biosciences, San Jose, CA) with appropriate excitation lasers and emission filters (Cerulean ex: 405 nm, em: 450/50 nm; Venus ex: 488 nm, em: 530/30 nm; FRET signals ex: 405 nm, em: 525/50 nm). Untransfected HEK293T cells served as gating controls to distinguish live, single-cell populations, and multiple single-channel controls (mCerulean-only, cpVenus-only, mCerulean and cpVenus co-transfection, AKAR6, and AKAR6 with Fsk/IBMX treatment) were included for accurate compensation and gating. Prior to measurement, cells were equilibrated for 10 minutes at room temperature. To analyze kinetics, continuous recordings were performed initially for 2 minutes at a low acquisition rate. Following baseline measurements, Fsk/IBMX (50 µM and 100 µM) were added, vortexed, and then recorded continuously for an additional 5 minutes. After this period, recording was briefly paused, and H89 (20 µM) was added, followed by vortexing and an additional 7-minute continuous recording.

Data were normalized by calculating the ratio of FRET to cerulean intensities. FlowJo software (v10.8.1, BD Biosciences) was used for data visualization and analysis.

### FLIM-FRET imaging

HeLa cells were imaged using a Leica SP8 FALCON confocal microscope (at the UCSD Microscopy Core) equipped with a 40×/1.3 NA oil-immersion objective, a white-light laser (WLL), a 440 nm pulsed diode laser (Picoquant, PDL-800-D) and a HyD SMD single molecule detector (Leica). For fluorescence imaging, mCerulean was excited at 440 nm and collected at 460–485 nm and cpVenus fluorescence was exited at 514 nm and collected at 525–555 nm. For FLIM imaging, mCerulean was excited at 440 nm at a repetition rate of 40 MHz. Time-course imaging was performed for 7 time points with 2-minute interval time, collecting photons for 30 s at each time point. Fsk/IBMX was added to cells after the second acquisition. A region of interest (ROI) corresponding to the cell cytosol was selected per cell. Fast FLIM maps (512×512 pixels), tail fitting of fluorescence decay curves, and amplitude-weighted mean fluorescence lifetimes were calculated in LAS X (Lecia) software by fitting a bi-exponential decay model to the decay of each ROI.

### Hippocampal slice culture and transfection

Animal work was performed in accordance with the Guide for the Care and Use of Laboratory Animals (by the US National Research Council Institute for Laboratory Animal Research), and was approved by the Institutional Animal Care and Use Committee (IACUC) of the Oregon Health & Science University (#IP00002274). Rat hippocampi were dissected from P6–7 pups of both sexes. Four-hundred-micron sections were prepared using a chopper. The dissection medium containing (in mM) 1 CaCl_2_, 5 MgCl_2_, 10 glucose, 4 KCl, 26 NaHCO_3_ and 248 sucrose, with the addition of 0.00025% phenol red. The slices were seeded onto a membrane (Millipore, PICM0RG50) and cultured at 35 °C with 5% CO2 in 7.4 g/l MEM (Thermo Fisher Scientific, 11700-077) plus 16.2 NaCl, 2.5 L-Glutamax, 0.58 CaCl_2_, 2 MgSO_4_, 12.9 D-glucose, 5.2 NaHCO_3_, 30 HEPES, 0.075% ascorbic acid, 1 mg/ml insulin and 20% heat-inactivated horse serum. Slice media were refreshed every 2–3 days by replacing ∼60% of the media. The cultured slices were transfected using the biolistic method (Bio-Rad Helios gene gun) at 10–14 days *in vitro* using 1.6 μm gold particles (Bio-Rad, 165-2262) coated with plasmids expressing the appropriate constructs (∼1 μg DNA per mg of gold). Cultured slices were imaged 2–3 days after transfection in carbogen-gassed aCSF containing (in mM) 126 NaCl, 26 NaHCO_3_, 10 D-glucose, 2.5 KCl, 1.25 NaH_2_PO_4_, 4 CaCl_2_, and 4 MgCl_2_.

### Two-photon imaging

The *ex vivo* two-photon microscope was built as previously described^42^ and controlled by the ScanImage software (Vidrio). mCerulean in both AKAR6 and AKAR6-H were excited with an 80 MHz pulsed titanium–sapphire laser at 850 nm. The fluorescence emission was unmixed using a Semrock FF511-Di01 dichroic mirror and Semrock FF01-483/32 barrier filter for mCerulean (donor channel) and Semrock FF01-550/49 barrier filter for cpVenus (FRET channel), respectively. Two-photon image analysis was carried out using previously described software suites called synScore^43^ written in MATLAB.

### Statistics and reproducibility

The number of cells analyzed (n cell) and number of independent experiments are reported in all figure legends. All replication attempts were successful. Statistical analyses were performed using GraphPad Prism 8 (GraphPad Software). For Gaussian data, pairwise comparisons were performed using Student’s t-test or Welch’s unequal variance t-test, and comparisons among three or more groups were performed using ordinary one-way analysis of variance followed by Dunnett’s test for multiple comparisons. Statistical significance was set at P□<□0.05.

