## Supplemental Figures for "Sensitive and Selective Next-Generation FRET-based PKA Biosensors"


**Figure S1**

**Figure S2**

**Figure S3**

**Figure S4**

**Figure S5**

**Figure S6**

**Figure S7**

**Figure S8**

**Figure S9**

**Figure S10**

**Figure S11**

**Figure S12**

**Figure S13**

**Table S1**

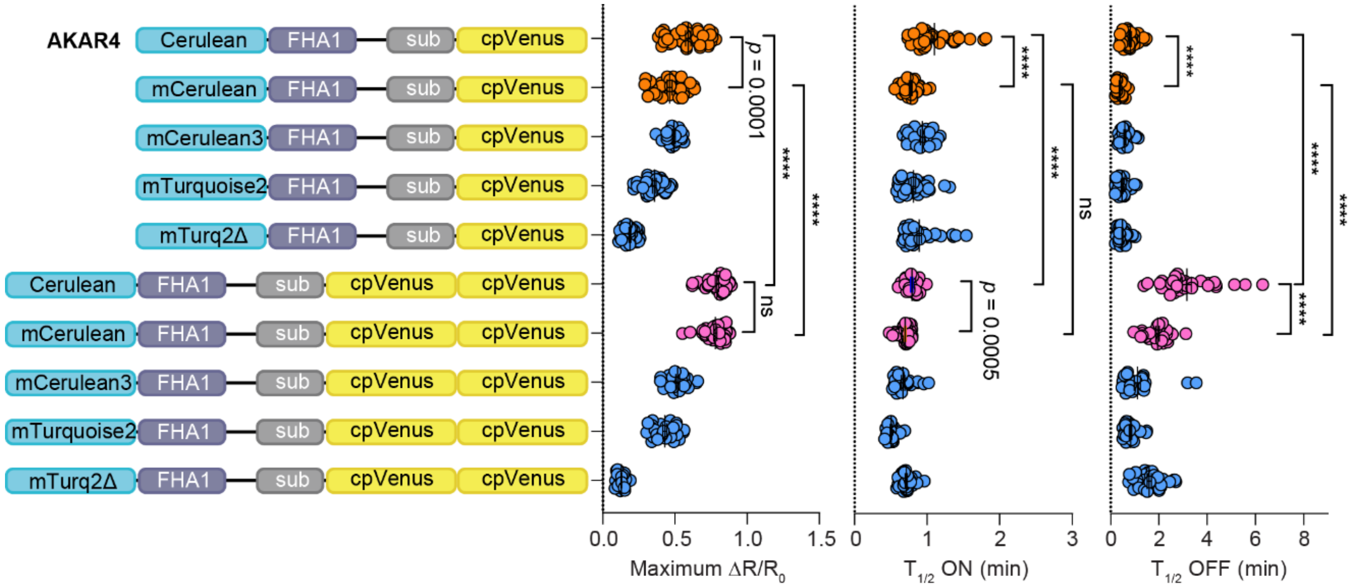


**Figure S1** Characterization of AKAR with different cyan-yellow FRET pairs. HeLa cell expressing AKARs with different cyan-yellow FRET pairs were treated with 50 μM Fsk and 100 μM IBMX for 5 min and 20 μM H89 for 10 min. Left panel: domain structures of AKAR variants. Right panel, from left to right: maximum responses ($\Delta R/R_0$), time to half-maximal activation ($T_{1/2}$ ON) and half-maximal inhibition ($T_{1/2}$ OFF). From top to bottom: n = 40, 33, 41, 28, 37, 34, 38, 37, 28 and 28 cells from 3 independent experiments each. Data indicate mean ± s.e.m. ****$P < 0.0001$; ns, not significant; Welch's ANOVA followed by Dunnett's test for multiple comparisons.

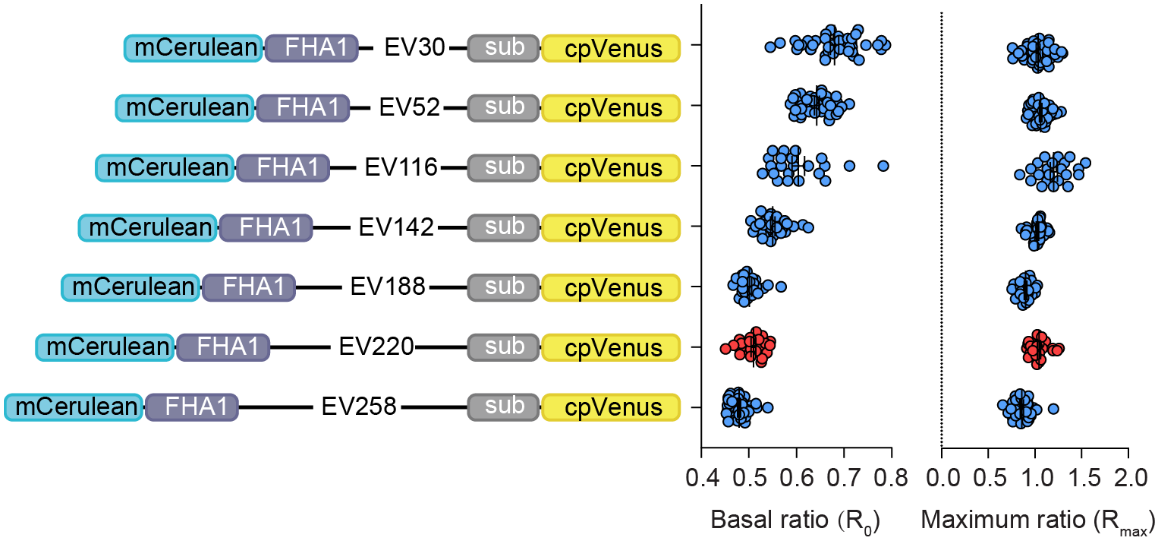


**Figure S2** Characterization of basal and maximal yellow/cyan emission ratio of mCerulean-cpVenus paired AKARs with different lengths of EV linkers. The EV (220 aa) variant (red) shows the largest normalized response ($\Delta R/R_0$) to Fsk/IBMX treatment. Left panel: domain structures of variants. Right panel, from left to right column: basal and maximum yellow/cyan emission ratios. From top to bottom: n = 41, 43, 22, 33, 26, 26, and 38 cells from three independent experiments each. Data indicate mean ± s.e.m.

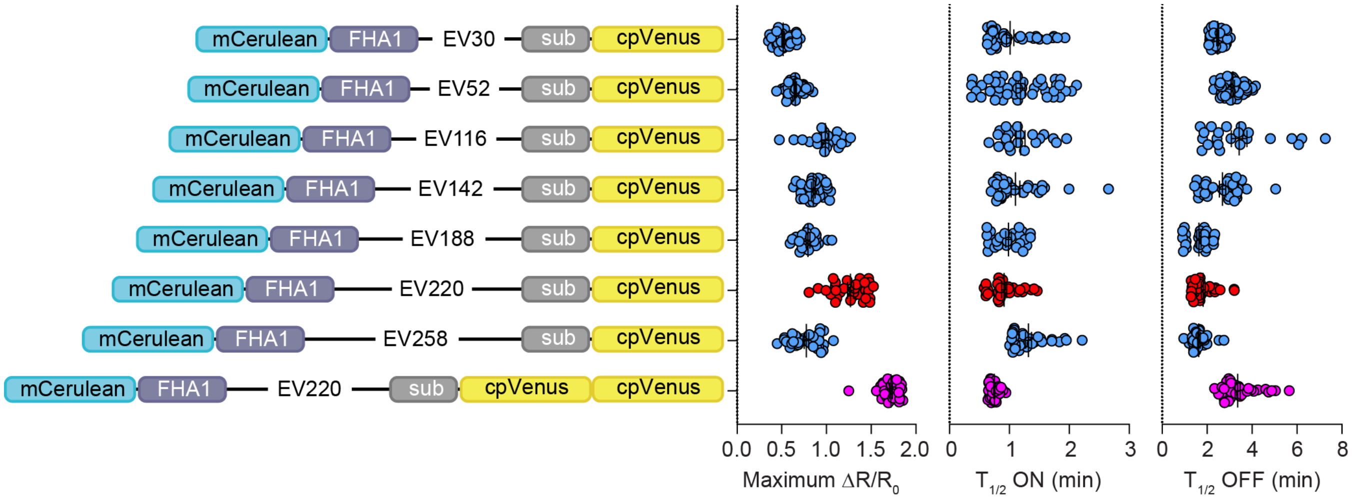


**Figure S3** Characterization of AKARs with different EV linker lengths. HeLa cell expressing mCerulean-cpVenus AKAR variants with different EV linkers lengths were treated with 50 µM Fsk and 100 µM IBMX for 5 min and 20 µM H89 for 10 min. Left panel: domain structures of variants. Right panel, from left to right column: maximum responses ($\Delta R/R_0$), time to half-maximal activation ($T_{1/2}$ ON) and half-maximal inhibition ($T_{1/2}$ OFF). From top to bottom: n = 39, 43, 22, 33, 26, 32, 38 and 44 cells from 3 independent experiments each. Data indicate mean ± s.e.m.

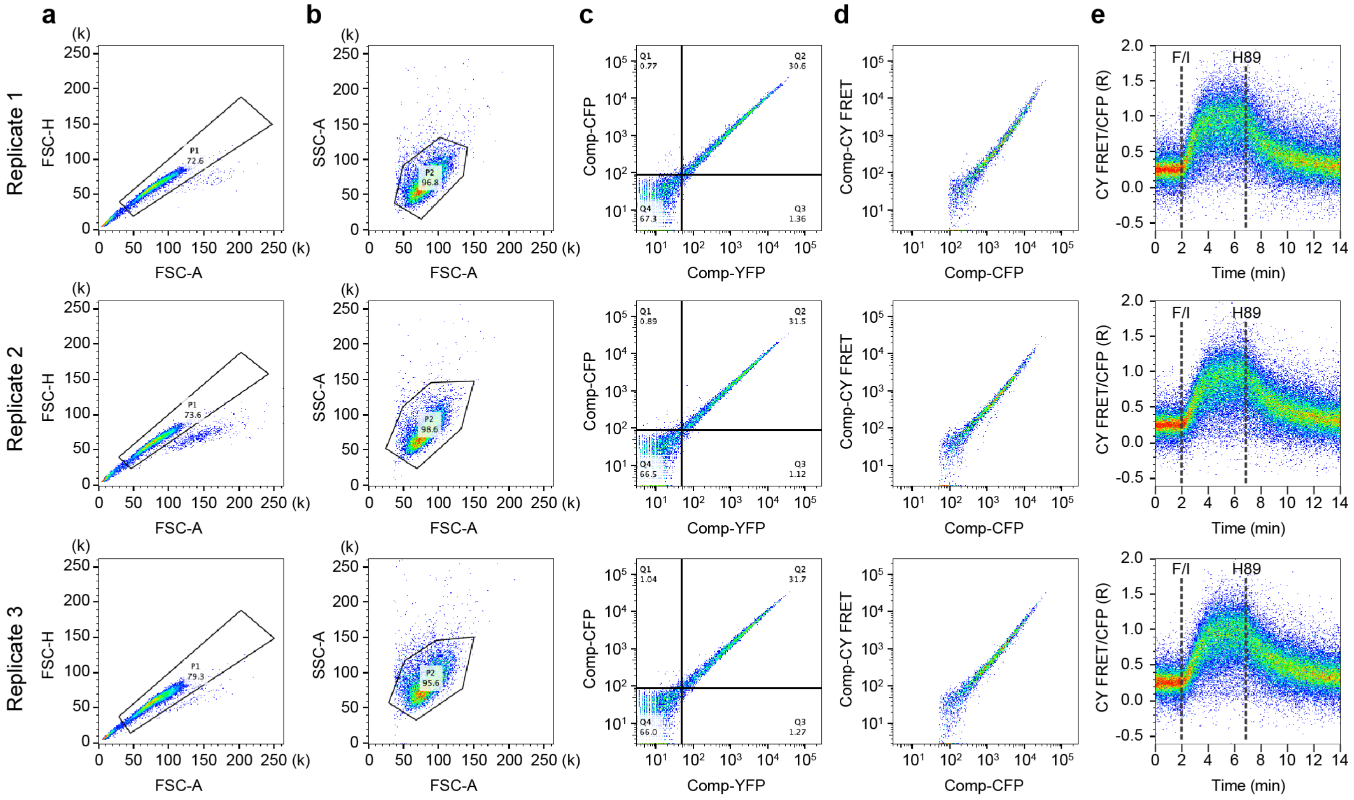


**Figure S4** Gating strategy for time-course ratiometric detection of AKAR6 response using flow cytometry. (a) Gating single cells. (b) Removing cell debris. (c) Gating CFP and YFP dual-positive cells (Q2). (d) The distribution of CY FRET vs CFP. (e) The scatter plots showing the ratio of CY FRET/CFP vs time course, during which Fsk/IBMX and H89 were sequentially added. The fluorescent intensity from each channel was corrected by compensation.

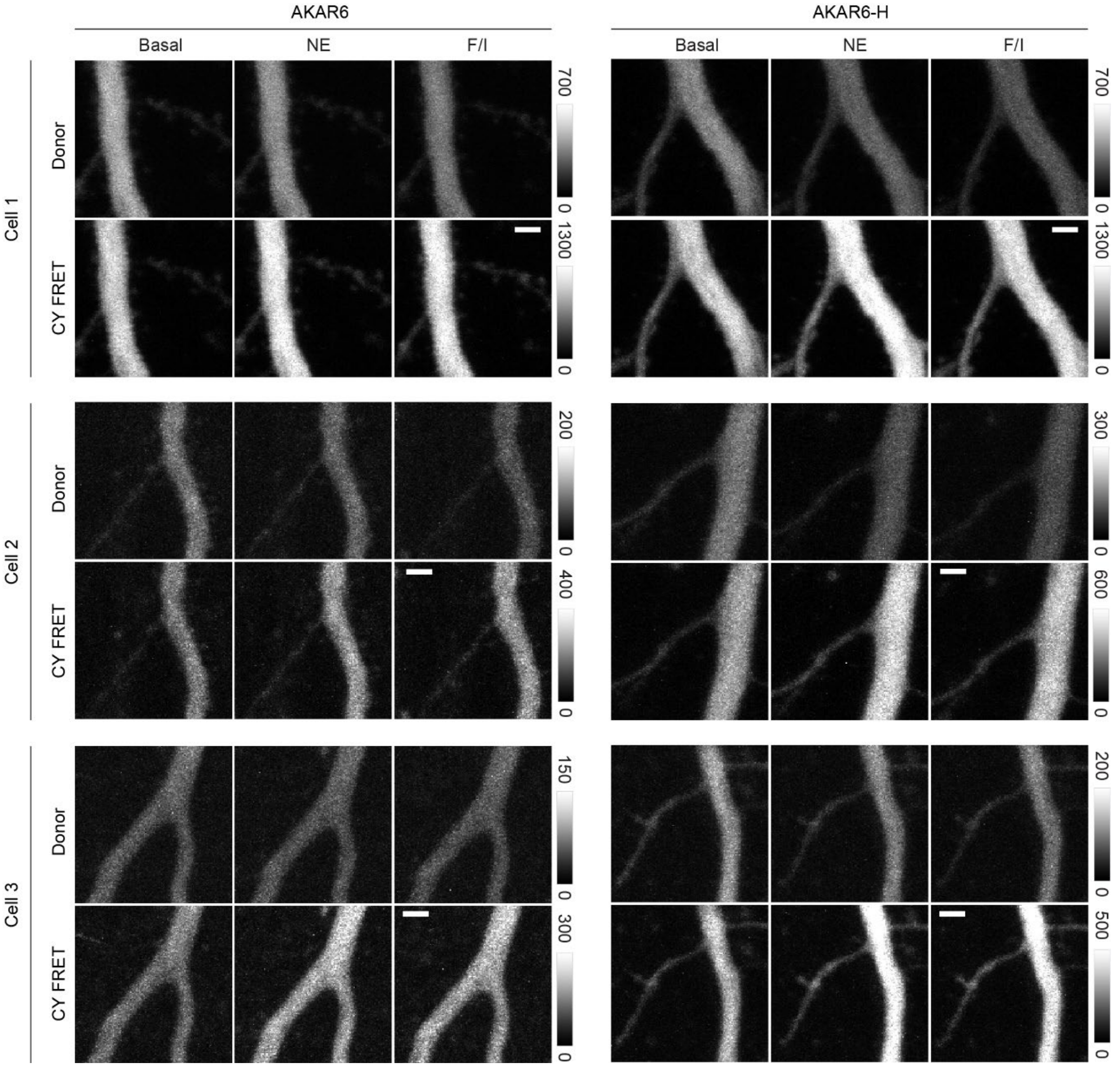


**Figure S5** Two-photon images of AKAR6 and AKAR6-H expressed in CA1 neurons of cultured hippocampal slices. Three CA1 neurons from three independent hippocampal slices for each sensor were sequentially treated with norepinephrine (NE) and Fsk/IBMX (F/I). Fluorescence images in donor and FRET channels were captured. Scale bars, 2 μm.

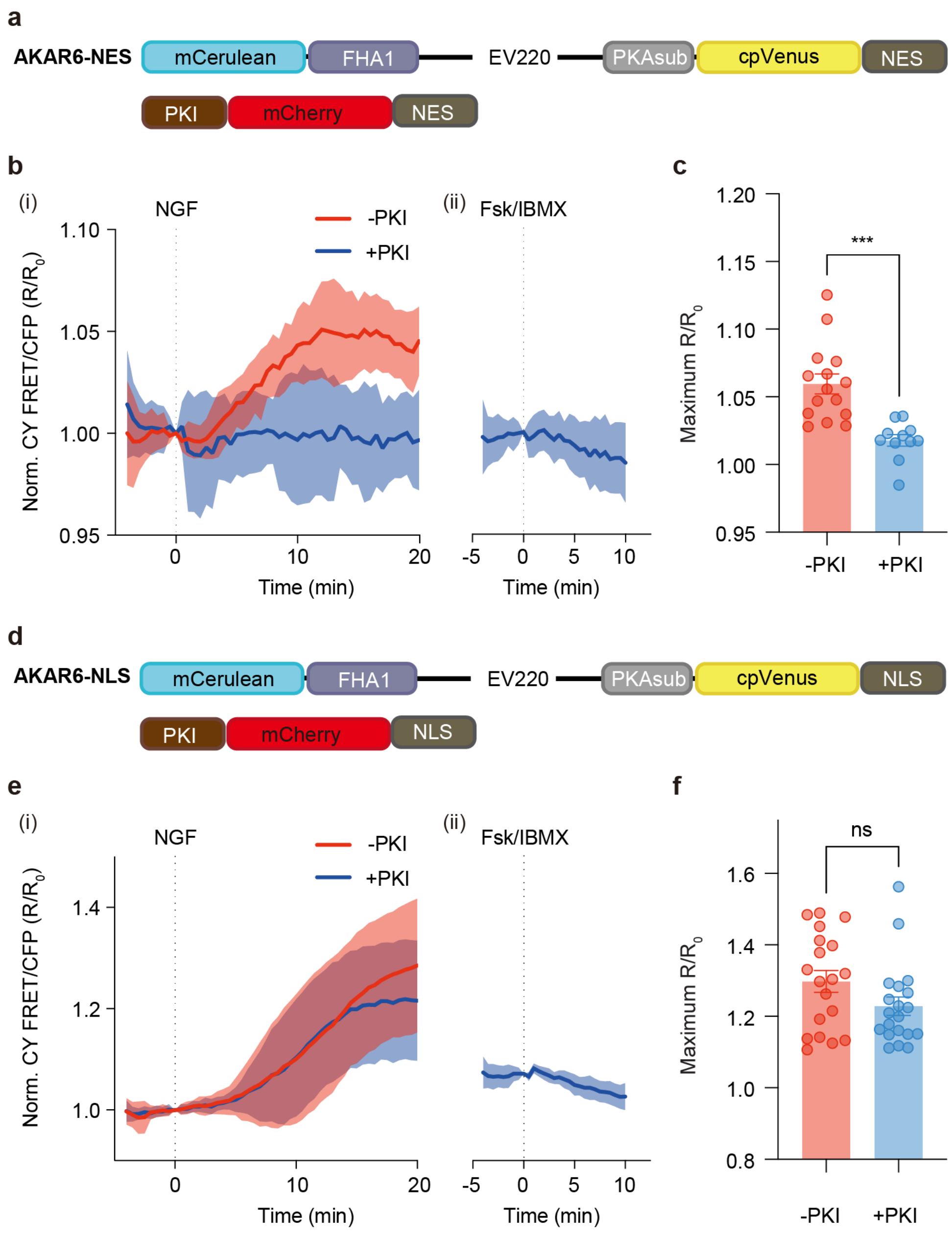


**Figure S6** Identification of non-specific nuclear AKAR6 phosphorylation in PC12 cells. (a) Domain structures of cytosol-targeted AKAR6 (AKAR6-NES) and PKI (PKI-mCherry-NES). (b) PC12 cells expressing AKAR6-NES alone respond to NGF stimulation (i, red curve), and co-expressing PKI-mCherry-NES abolishes the response to both (i) NGF and (ii) Fsk/IBMX stimulation (blue curves). (NGF, -PKI, n = 15 cells; NGF, +PKI, n = 11 cells; Fsk/IBMX, +PKI, n = 12 cells). (c) Quantification of maximum NGF-stimulated AKAR6-NES responses ($R/R_0$). (d) Domain structures of nuclear-targeted AKAR6 (AKAR6-NLS) and PKA (PKI-mCherry-NLS). (e) AKAR6-NLS-expressing PC12 cells respond to NGF stimulation both without (red curve) and with (blue curve) PKI-mCherry-NLS co-expression (i), whereas PKI-mCherry-NLS abolishes the response to (ii) Fsk/IBMX stimulation, indicating that NGF induces non-specific AKAR6 phosphorylation in the nucleus. (NGF, -PKI, n = 19 cells; NGF, +PKI, n = 20 cells; Fsk/IBMX, +PKI, n = 12 cells). (f) Quantification of maximum NGF-stimulated AKAR6-NLS responses

($R/R_0$). Solid lines in b and d indicate mean responses and shaded areas indicate s.d. Data in c and f are mean ± s.e.m. ***$P < 0.001$; ns, not significant; unpaired two-tailed Student's *t*-test.

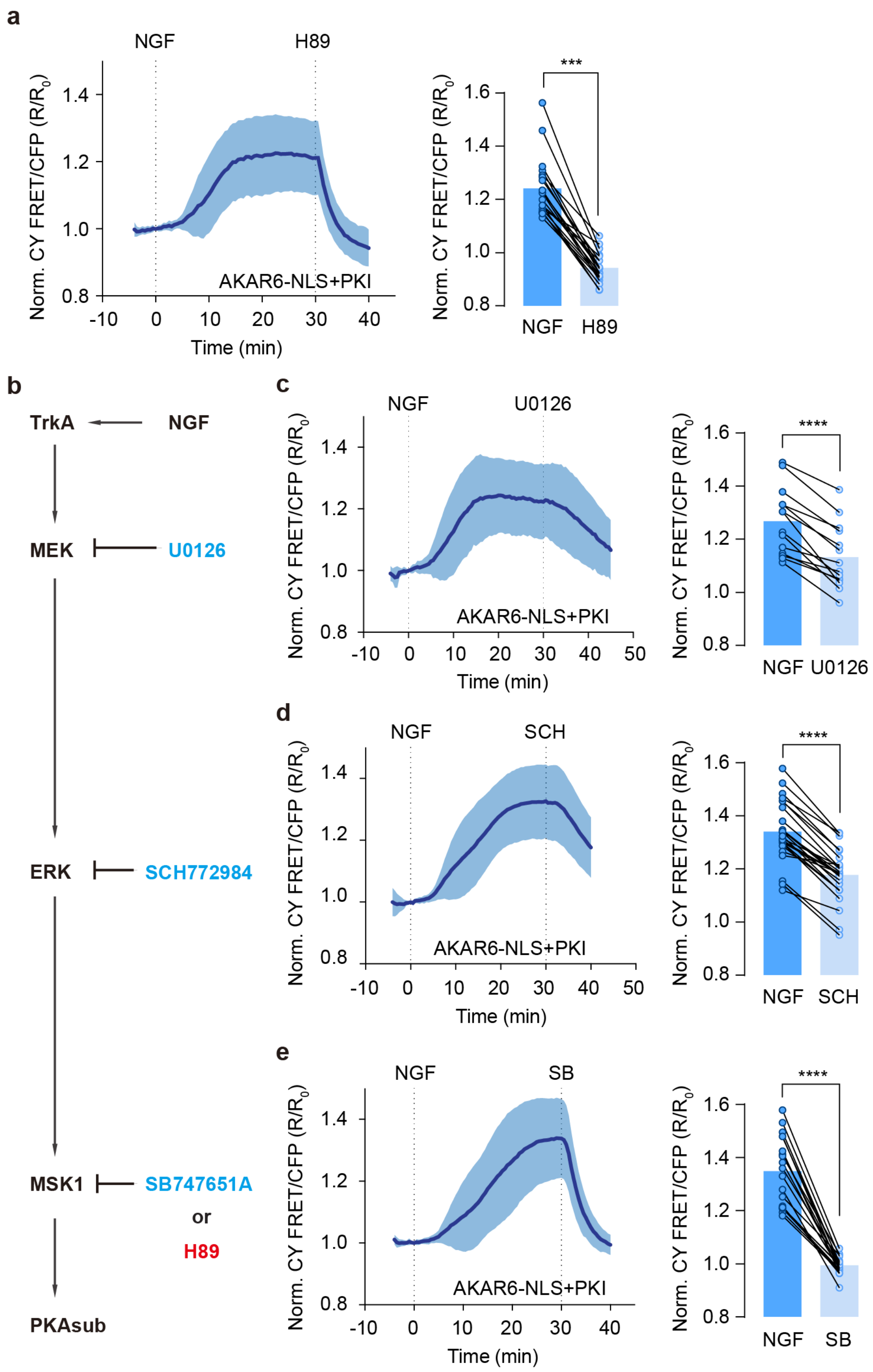


**Figure S7** Identification of MSK1 phosphorylation on original PKA substrate. (a) Representative time-course responses of AKAR6-NLS to NGF-induced nuclear non-PKA activity which can be non-specifically inhibited byH89 and quantification of maximum responses ($R/R_0$) to NGF and final response ($R/R_0$) to H89 inhibition (n = 21 cells). (b) Schemes of the MEK-ERK-MSK1 pathway and phosphorylation of original PKA substrate. Arrowheads indicate direct phosphorylation and blunt arrowheads indicate inhibition. H89 is an ATP analogue inhibiting multiple kinases including MSK1 (red). (c-e) NGF induced AKAR6-NLS responses can be inhibited by U0126 (c), SCH772984 (d) and SB747651A (e). From top to bottom, n = 14, 25 and 19 cells. Data from 3 independent experiments. Solid lines indicate mean responses and shaded areas indicate the s.d. ***$P < 0.001$, ****$P < 0.0001$. Data were analyzed using paired two-tailed Student's t test. Data are mean ± s.e.m.

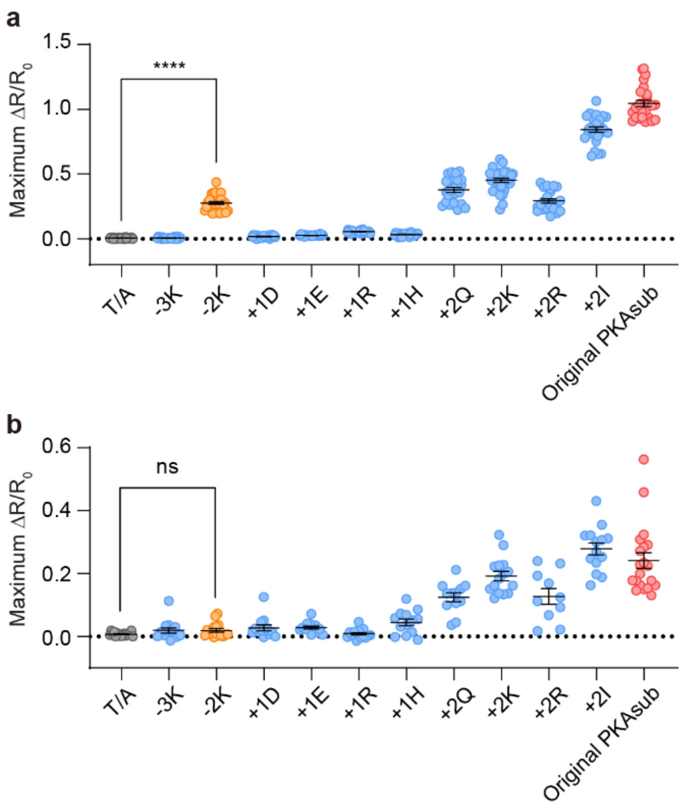


**Figure S8** Characterization of PKA and MSK1 phosphorylation on different PKA substrate mutants. Phosphorylation is reflected by maximal normalized response ($\Delta R_{max}/R_0$) of AKAR6 substrate mutants under PKA and MSK1 specific phosphorylation. AKAR6 with the original PKA substrate can be phosphorylated by both PKA and MSK1 (red, while the mutant -2K can only be phosphorylated by PKA (orange). AKAR6 T/A serves as a negative control that cannot be phosphorylated (black). (a) AKAR6 with substrate mutants are expressed in HeLa cells. Fsk/IBMX induce maximal PKA activity to phosphorylate the sensor. From left to right: n = 27, 21, 30, 22, 21, 28, 26, 25, 30, 24, 26 and 26 cells from 3 independent experiments each. ****$P < 0.0001$. Data were analyzed using Welch's ANOVA followed by Dunnett's test for multiple comparisons to T/A. (b) AKAR6-NLS with substrate mutants are expressed in PC12 cells, and PKI-mCherry-NLS is co-expressed to inhibit nuclear PKA activity. NGF induces strong activation of MSK1 activity in nucleus. From left to right: n = 17, 14, 16, 13, 14, 17, 13, 12, 15, 10, 14 and 21 cells from 3 independent experiments each. ns, not significant. Data were analyzed using Welch's ANOVA followed by Dunnett's test for multiple comparisons to T/A.

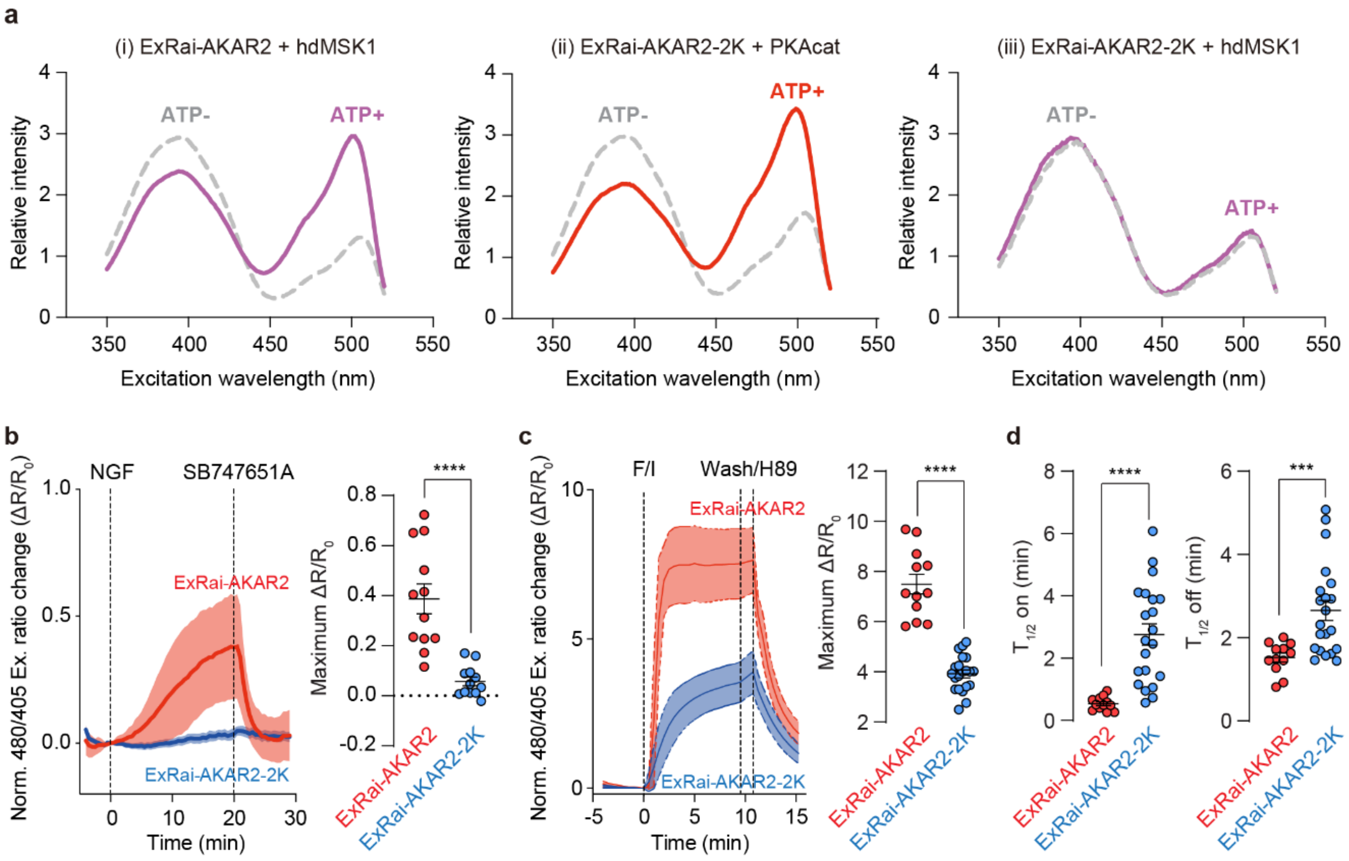


**Figure S9** The "-2K" mutation improves the kinase selectivity of ExRai-AKAR2. (a) Representative ExRai-AKAR2 and ExRai-AKAR2 -2K mutant fluorescence spectra collected at 530 nm emission. ExRai-AKAR2 is incubated with truncated human MSK1 (i), ExRai-AKAR2 -2K is incubated with PKA catalytic subunit (ii) and truncated human MSK1 (iii), respectively, without (gray dashed curve) or with (red or magenta solid curve) ATP. (b) Representative time-course responses of ExRai-AKAR2 (red, n = 12 cells) and ExRai-AKAR2-2K (blue, n = 12 cells) to NGF stimulation and MSK1 inhibition in PC12 cell (left). Quantification of maximum responses ($\Delta R/R_0$) of ExRai-AKAR2 and ExRai-AKAR2-2K (right). ****$P < 0.0001$. Data were analyzed using unpaired two-tailed Student's *t*-test with Welch's correction, (c) Representative time-course responses of ExRai-AKAR2 (red, n = 12 cells) and ExRai-AKAR2-2K (blue, n = 21 cells) in HeLa cell treated with Fsk/IBMX for 10 min and 20 μM H89 for 5 min (left). Quantification of ExRai-AKAR2 and ExRai-AKAR2-2K maximum responses ($\Delta R/R_0$) to Fsk/IBMX (right). Solid lines indicate mean responses, shaded areas indicate the s.d. ****$P < 0.0001$. Data were analyzed using unpaired two-tailed Student's *t*-test with Welch's correction. (d) Quantification of ExRai-AKAR2 and ExRai-AKAR2-2K time to half-maximal activation ($T_{1/2}$ ON) and half-maximal inhibition ($T_{1/2}$ OFF). ****$P < 0.0001$. Data were analyzed using unpaired two-tailed Student's *t*-test with Welch's correction. All data from 3 independent experiments each. Data are mean ± s.e.m.

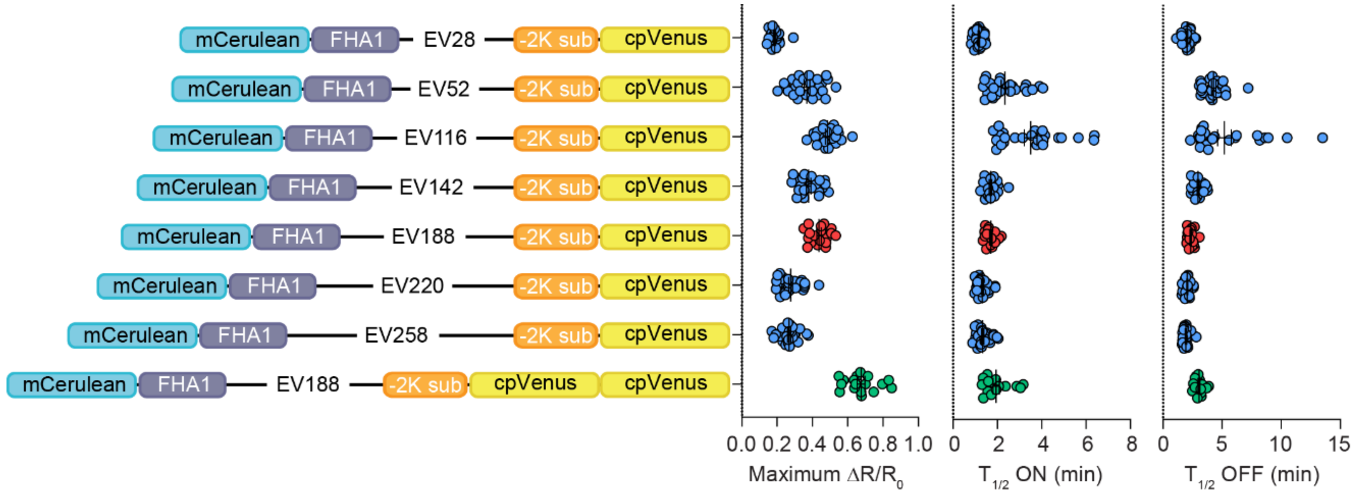


**Figure S10** Optimization of AKAR6(−2K). Left panel: Domain structures of AKAR6(−2K) and its variants. Right panel, from left to right column: Maximum responses ($\Delta R/R_0$), time to half-maximal activation ($T_{1/2}$ ON) and half-maximal inhibition ($T_{1/2}$ OFF). From top to bottom: n = 28, 26, 25, 24, 18, 30, 26 and 17 cells from three independent experiments each. The optimal linker is EV188 (red), leading to relatively large dynamic range and fast kinetics. This variant is further improved by switching to tdcpVenus (green) and designated as AKAR6-S.

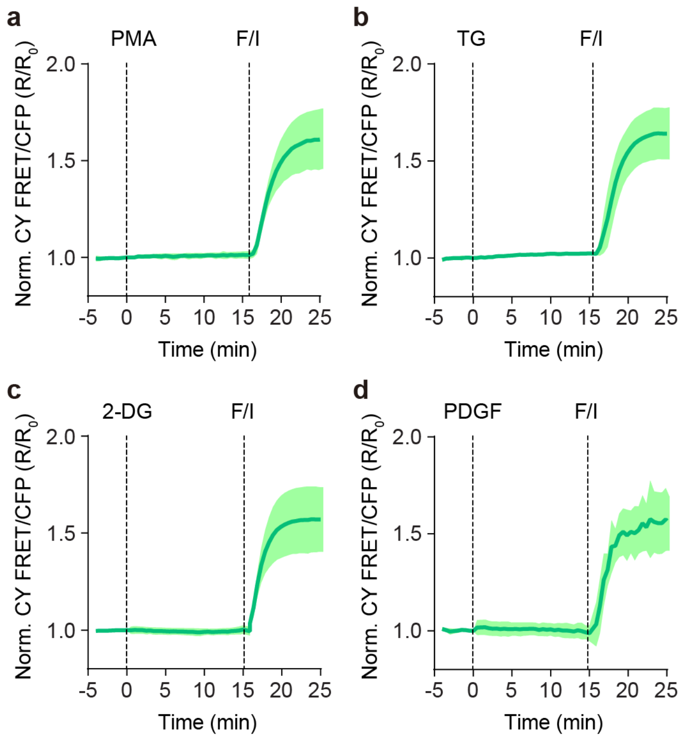


**Figure S11** AKAR6-S does not respond to PKC, CaMKII, AMPK and Akt stimulation. Representative time-course responses of AKAR6-S in (a-c) HEK293T cell stimulated with (a) 100 ng ml$^{-1}$ PMA to activate PKC (n = 24 cells), (b) 1 µM thapsigargin (TG) to activate CaMKII (n = 16 cells), (c) 40 mM 2-deoxyglucose to activate AMPK (n = 17 from 3 experiments) and (d) in serum-starved NIH3T3 cell stimulated with 50 ng mL$^{-1}$ of PDGF to activate AKT (n = 11 cells). Cells were treated with 50 µM Fsk and 100 µM IBMX at 15 min as a positive control. Data are representative of 3 independent experiment each. Solid lines indicate mean responses; shaded areas, s.d.

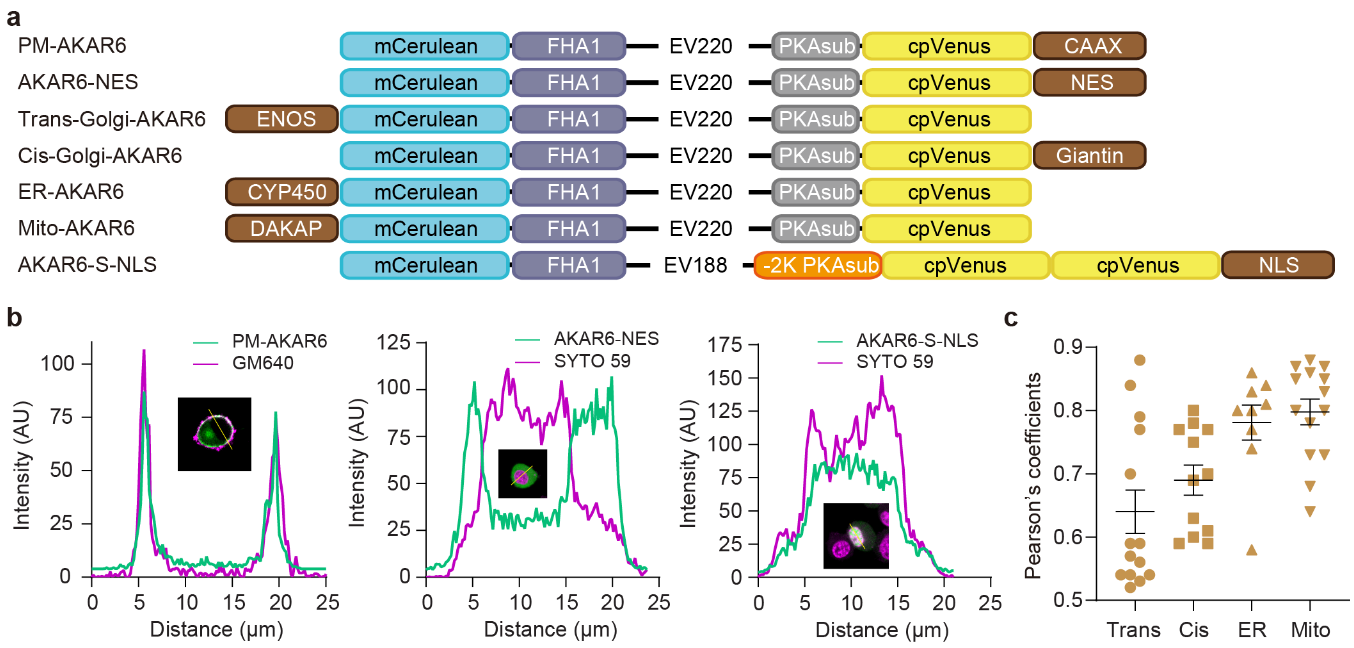


**Figure S12** Subcellularly targeted AKAR6 and AKAR6-S. (a) Domain structures of subcellular localized AKAR6 and AKAR6-S. Specific subcellular markers including CAAX, NES, ENOS, Giantin, CYP450, DAKAP and NLS are added to N- or C-terminal of sensors to generate PM-AKAR6, AKAR6-NES, Trans-Golgi-AKAR6, Cis-Golgi-AKAR6, ER-AKAR6, Mito-AKAR6 and AKAR6-S-NLS, respectively. (b) Intensity profiles of representative dual-color fluorescence images showing subcellular AKAR6 variants with markers (left, PM-AKAR6 merged with plasma membrane staining dye GM640, middle, AKAR6-NES merged with DNA staining dye SYTO 59, right, AKAR6-S-NLS merged with DNA staining dye SYTO 59). (c) Quantitative analysis using Pearson's coefficient indicates colocalization of Trans-Golgi-AKAR6, Cis-Golgi-AKAR6, ER-AKAR6 and Mito-AKAR6 with corresponding markers (n = 14, 12, 9 and 14 cells) from 3 independent experiments each.

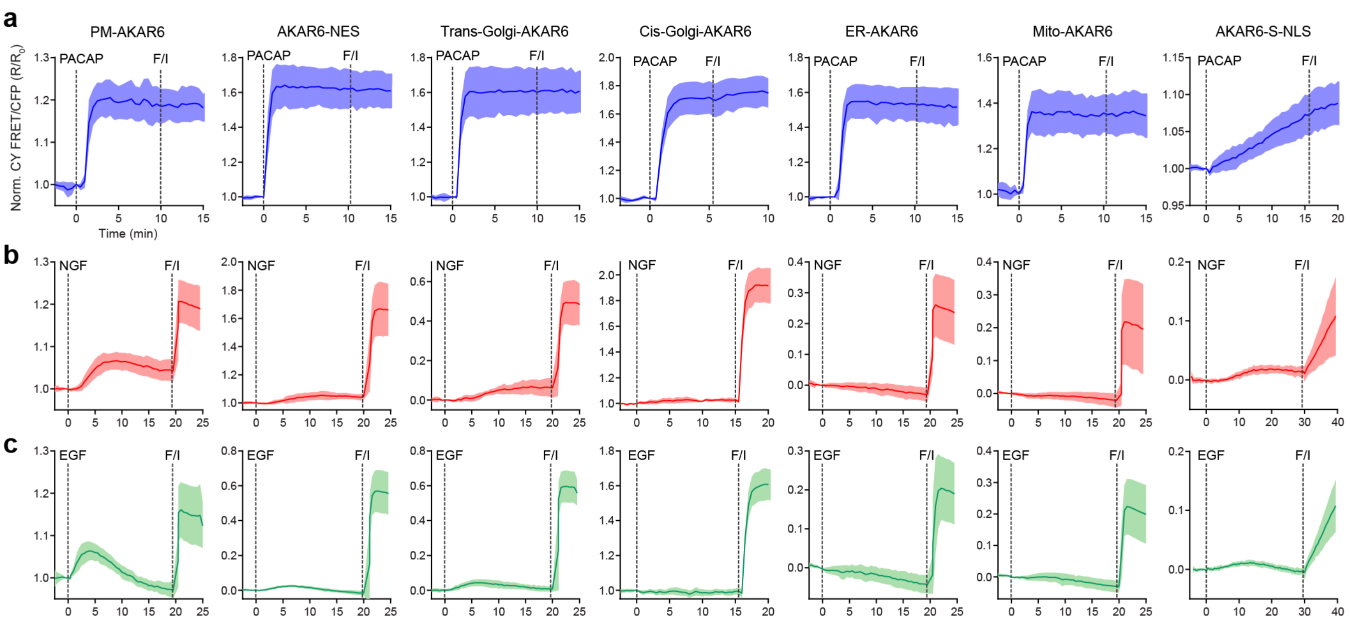


**Figure S13** Full time-courses of seven subcellular targeted AKAR6 variants responding to PACAP, NGF or EGF stimulations followed by Fsk/IBMX. Blue (a), red (b) and green (c) curves indicate PKA activities stimulated with 100 nM PACAP, 200 ng $mL^{-1}$ NGF and 100 ng $mL^{-1}$ EGF, respectively. Solid lines indicate mean responses; shaded areas, s.d. (PM-AKAR6, PACAP, n = 5 cells, NGF, n = 11 cells, EGF, n= 7 cells; AKAR6-NES, PACAP, n = 6 cells, NGF, n = 10 cells, EGF, n = 12 cells; Trans-Golgi-AKAR6, PACAP, n = 9 cells, NGF, n = 11 cells, EGF, n = 14 cells; Cis-Golgi-AKAR6, PACAP, n = 6 cells, NGF, n = 13 cells, EGF, n = 17 cells; ER-AKAR6, PACAP, n = 6 cells, NGF, n = 12 cells, EGF n = 12 cells; Mito-AKAR6, PACAP, n = 7 cells, NGF, n = 17 cells, EGF, n = 12 cells; AKAR6-S-NLS, PACAP, n = 6 cells, NGF, n = 11 cells, EGF, n = 12 cells). Data representative of 3 independent experiments each.

**Table S1 Primers used for molecular cloning**

| Primer Number | Name | sequence (5' to 3') |
|---|---|---|
| 1 | EFP-BamHI_F | GGCGGATCCCATGGTGAGCAAGGGCGAG |
| 2 | EC-SphI_R | GGCGATGCATGCGGGCGGCGGTCAC |
| 3 | PKAsub_Sac1_cpV172_F | CTGGTGGACGGCGGCACCGGCGGCAGCGAGCTCATGGACGGCGGCGTG |
| 4 | td-cpV-linker_R | GCTAGATACGAAGAGTATTCCTGCGGCGCCGATCAGGAAGAC |
| 5 | td-cpV-linker_Kpn1_F | CCGAATTTTGTCTTCCTGATCGGCGCCGCAGGAATACTCTTCGTATCTAGCGGTACCATGGACGGCGGCGTG |
| 6 | pRSET_EcoR1_stop_cpV172_R | CGGGCTTTGTTAGCAGCCGGATCAAGCTTCGAATTCTTACTCGATGTTGTGGCG |
| 7 | pc3prime-Bam-FP_F | GGTCGGGATCTGTACGACGATGACGATAAGGATCCCATGGTGAGCAAGGGCGAGGAG |
| 8 | FHA1-Sph-FP_R | GCCGATCTGTTCTTGAGAAAACTTATGCATGCGCTTGTACAGCTCGTCCATGCC |
| 9 | Cerulean_A206K_F | CACTACCTGAGCACCCAGTCCAAGCTGAGCAAAGACCCCAACGAG |
| 10 | Cerulean_A206K_R | CTCGTTGGGGTCTTTGCTCAGCTTGGACTGGGTGCTCAGGTAGTG |
| 11 | backbone-split_F | ATTCTAGTTGTGGTTTGTCCAAACTCATCAA |
| 12 | backbone-split_R | TTGATGAGTTTGGACAAACCACAACTAGAAT |
| 13 | EV-Kpn-FHA_R | ACTACCACCAGCACTACCACCAGCACTGGTACCGCGATCAACTTTGTTCTGCTCGAGGCA |
| 14 | EV-BspE-PKAsub_F | GGTGGTAGTGCTGGTGGTAGTGCTGGTGGTTCCGGACTGCGTCGCGCCACCCTGGTGGAC |
| 15 | EV- Kpn-FHA_F | GCAGAACAAAGTTGATCGCGGTACCAGTGCTGGTGGTAGTGCTGGTGGTAGTGCTGGTGGTAGTGCTGGTGGTAGTGCTGGTGGTTCCGGCAGTGCTGGTGGTAGTGCTGGTGGTAGTACCAGTGCTGGTGGTAGTGCTG |
| 16 | EV-BspE-PKAsub_R | ACCAGGGTGGCGCGACGCAGTCCGGAACCACCAGCACTACC |
| 17 | PKAsub_+3D_F | GACGGCGGCACCG |
| 18 | PKAsub_-3K_R | CTGCCGCCGGTGCCGCCGTCCACCAGGGTGGCGCGCTTCAGTCCGGAACCACCAGC |
| 19 | PKAsub_-2K_R | CTGCCGCCGGTGCCGCCGTCCACCAGGGTGGCAAGACGCAGTCCGGAACCACCAGC |
| 20 | PKAsub_+1R_R | CTGCCGCCGGTGCCGCCGTCCACGCGGGTGGCGCGACG |
| 21 | PKAsub_+1D_R | CTGCCGCCGGTGCCGCCGTCCACGTCGGTGGCGCGACGC |

| | | |
|---|---|---|
| 22 | PKAsub_+1E_R | CTGCCGCCGGTGCCGCCGTCCACCTCGGTGGCGCGACGC |
| 23 | PKAsub_+1H_R | CTGCCGCCGGTGCCGCCGTCCACGTGGGTGGCGCGACG |
| 24 | PKAsub_+2Q_R | CTGCCGCCGGTGCCGCCGTCCTGCAGGGTGGCGCGAC |
| 25 | PKAsub_+2K_R | CTGCCGCCGGTGCCGCCGTCCTTCAGGGTGGCGCGAC |
| 26 | PKAsub_+2R_R | CTGCCGCCGGTGCCGCCGTCACGCAGGGTGGCGCGAC |
| 27 | PKAsub_+2I_R | CTGCCGCCGGTGCCGCCGTCGATCAGGGTGGCGCGAC |
| 28 | EV_BspEI_PKAs ub_Gibson_F | GTGGTAGTGCTGGTGGTTCCGGACTGCGTCGCGCCA |
| 29 | cpVenus_tdlinker _R | GAGTATTCCTGCGGCGCCGATCAGGAAGACAAAATTCG GCTCGATGTTGTGGCGGATCTT |
| 230 | tdLinker_ cpVenus_F | ATCGGCGCCGCAGGAATACTCTTCGTATCTAGCGGTACC ATGGGCGGCGTGC |
| 31 | PCDNA3_XbaI_R | ACTATAGAATAGGGCCCTCTAGA |
